# Nitrogen fertilization outweighs plant species loss in shaping bacterial belowground diversity in an alpine meadow on the central Tibetan Plateau

**DOI:** 10.64898/2026.04.08.717155

**Authors:** Dongmei Wu, CiRen Quzong, Zhongjun Jia, Antje Schwalb, Georg Guggenberger, Shipeng Wang, Tsechoe Dorji, Michael Pester

## Abstract

Plant species loss and nitrogen fertilization affect grassland biodiversity. However, their interactive effects on plant communities, soil properties, and the soil microbiome remain insufficiently understood. We analyzed how the removal of plant species, with and without urea addition, influenced plant diversity, soil properties, and soil bacterial communities in a Tibetan Plateau grassland. Continuous plant species removal and urea addition over seven years modified plant beta-diversity equally strong, while urea exerted a stronger negative effect on plant alpha-diversity. Both, plant species removal and urea addition caused soil acidification and an increase in NO₂⁻/NO₃⁻, while dynamics in TOC, TON and TOC: TON were mainly driven by the growing season. Structural equation modeling identified soil acidification via urea addition as the most important indirect driver that negatively affected bacterial alpha-diversity and shifted bacterial beta-diversity. Urea addition also exerted direct negative effects on bacterial alpha- and beta-diversity, causing repression of oligotrophic (*Acidobacteriota*, *Chloroflexota*, *Planctomycetota*, *Gemmatimonadota*) and stimulation of copiotrophic (*Bacillota*, *Bacteroidota*, *Pseudomonadota*) bacterial taxa. Plant species removal caused slight increases in bacterial alpha-diversity, paralleled by less diverse but more even plant communities. We show that soil acidification by urea fertilization outweighs plant species loss in its negative effect on bacterial soil biodiversity in Tibetan grasslands.

## Introduction

Plant-soil feedbacks describe the complex interactions between plants, soil properties and (micro-)organisms inhabiting the soil, which feedback on plant performance, diversity and community structure, ultimately resulting in ecosystem functioning and stability (Van der Putten et al. 2013, Pugnaire et al. 2019, Cheng et al. 2025). Dominant plant species play a significant role in these feedback mechanisms (Wardle et al. 1999, Chaves and Smith 2021, Hernández et al. 2022, Rewcastle et al. 2022, Wang and Li 2024) through root exudates, secondary metabolites and their influence on soil properties, such as pH or nutrient content. Therefore, they affect directly and indirectly the soil microbiome (Wardle et al. 2004, Bezemer et al. 2006, Eisenhauer et al. 2010, Ladygina and Hedlund 2010, Ward et al. 2015, Stefanowicz et al. 2018). These interactions help maintain soil health and nutrient cycling, making dominant plants key drivers in structuring the microbial community composition and its function within ecosystems (Chen et al. 2019, Beugnon et al. 2021, Chen and Chen 2021, Sokol et al. 2022, Philippot et al. 2024). When such species are lost, the composition and function of soil microbial communities can shift significantly, which can negatively feedback on ecosystem functionality and biodiversity (Knops et al. 2002, Eisenhauer et al. 2010, Xiao et al. 2017, Chen et al. 2019).

Anthropogenic N loading, e.g. by fertilizer application or atmospheric deposition (Fowler et al. 2013, Gu et al. 2015, Yu et al. 2019), is an important driver of global change that significantly alters soil properties, impacting plant and microbial community structure as well as nutrient cycling (Bobbink et al. 2010, Cleland and Harpole 2010, Ramirez et al. 2010, Chen et al. 2019, Han et al. 2019, Wang et al. 2023, Liao et al. 2024, Ma et al. 2025). In particular, nitrogen inputs concomitant with soil acidification can negatively affect soil microbial communities, often reducing microbial biomass and respiration rates (Wei et al. 2013, Tian et al. 2017, Zhang et al. 2018). This decline in microbial activity can disrupt essential soil functions, diminishing nutrient turnover and carbon cycling (Chen et al. 2015, Wang et al. 2025).

Both, plant species loss and anthropogenic N loading are relevant for the Tibetan Plateau, a climate-relevant region holding extensive C-rich alpine grasslands (Jin et al. 2005, Yao et al. 2012, Miehe et al. 2019). The latter store about 33.5 Pg C spread over 1.6 million km^2^ (Genxu et al. 2002, Yao et al. 2012), with the 450,000 km^2^ of *Kobresia pygmea*-dominated pastures alone contributing 21.7 Pg C to these soil organic carbon stocks (Miehe et al. 2019). This is especially fostered by the dwarf growth of *K. pygmea*. This sedge is the smallest of the High Asian *Cyperaceae*, growing no more than 2 cm tall and allocating most of its photoassimilates belowground (Miehe et al., 2008). The resulting large root biomass stores nutrients belowground, ensuring fast regrowth following grazing events to cover the high belowground C costs (Miehe et al. 2019). As a result, these grasslands sustain 13 million yaks and 30 million goats and sheep, which support the livelihood of ca. 5 million pastoralists (Miehe et al. 2019).

About 14-16% of the Tibetan grasslands are already degraded (Genxu et al. 2002, Yao et al. 2012). Nutrient-poor soils and adaptation to extreme climatic conditions (low temperature, wide diurnal temperature range, high UV radiation, low O_2_ and CO_2_ partial pressure), along with widespread endemism among dominant plant species, make this ecosystem particularly vulnerable to environmental changes (Miehe et al. 2008, Yao et al. 2012, Cheng et al. 2023, Dong et al. 2023). Atmospheric warming on the Tibetan Plateau occurs approximately 2-3 times faster than on the global average (Liu and Chen 2000), with an predicted increase of an additional 4°C within the next 100 years (Liu and Chen 2000, Yao et al. 2019). Such rapid temperature increases threaten native alpine plant species by altering the delicate environmental conditions they require for survival. For instance, warming has already led to declines in both medicinal and forage plant species, critical for local biodiversity and human use (Klein et al. 2008, Wang et al. 2012). It is estimated that, due to climate warming, plant species richness may decrease by >25% on the Tibetan Plateau (Klein et al. 2004, Li et al. 2024) with negative effects for soil quality (Zhou et al., 2019).

Strong local N loadings on the Tibetan Plateau may result from restoration efforts of degraded grasslands. A recent meta-analysis identified 272 kgLNLha^−1^Lyr^−1^ as an optimal N fertilization rate to restore degraded grasslands on the Qinghai-Tibetan Plateau (He et al. 2024). In addition, atmospheric N deposition has increased since the 1980s at an average rate of about 0.17LkgLNLha^−1^Lyr^−1^ reaching values of 4.0 kgLNLha^−1^Lyr^−1^ in the 2000s (Liu et al. 2013, Chen et al. 2022), with expectations of further increases (Gu et al. 2015). The lower input of atmospheric N deposition alone has already been shown to decrease plant species richness and functional diversity (Han et al. 2019, Shen et al. 2022, Xiang et al. 2025) as well as to influence soil microbial communities (Xiong et al. 2016, Zhang et al. 2020) of grasslands on the Tibetan Plateau, in turn impacting soil processes and greenhouse gas dynamics (Liu and Zhang 2019, Xu et al. 2021). While the effects of warming and increased N loading on grasslands on the Tibetan Plateau have been studied for plant (Xiang et al. 2025) and soil microbial communities (Xiong et al. 2016, Zhang et al. 2020, Mu et al. 2021) separately, the combined effects of dominant plant species loss and nitrogen fertilization on above- and belowground biodiversity remain largely unknown. To address this, we utilized a long-term experimental station involving selective plant species removal and addition of urea as a common fertilizer over a period of seven years to explore how the loss of dominant plant species and strong N fertilization affect soil microbial communities on the Tibetan Plateau. In particular, we tested (1) whether plant species loss or urea addition would affect soil bacterial diversity stronger and (2) whether both tested factors exert strong direct effects or rather act via indirect effects on soil bacterial diversity.

## Materials and Methods

### Study site and design

The studied alpine grassland (31°16’ N, 92°05’ E, 4,512 m above sea level) is part of the Nagqu Alpine Grassland Ecosystem National Field Scientific Observation and Research Station (hereafter, Nagqu station), run by the Tibet University and the Institute of Tibetan Plateau Research, Chinese Academy of Sciences. The area is characterized by intense solar radiation, long cold winters, and short, cool summers (Li et al. 2016). The average annual air temperature is −1.2°C, with an annual precipitation of 432 mm (Li et al. 2023), most of which occurs during the warm season from May to September (Miehe et al. 2008). The grassland is dominated by the endemic *Cyperaceae* (sedge) member *Kobresia pygmaea* C.B. Clarke (Table S1), which covers 75-98% of the total plant community in south-eastern Tibetan grasslands (Miehe et al. 2008). The *Rosaceae* member *Potentilla saundersiana L*. is another common plant species widely distributed across Tibetan alpine grasslands (https://powo.science.kew.org/) (Ma et al. 2015, Liu et al. 2021), covering up to 2-4% of the plant community (Miehe et al. 2008, Table S1). Both plant species differ in their flowering period, which is early in the growing season for *K. pygmaea* and mid-season for *P. saundersiana* (Suonan et al., 2024). In 2014, the grassland was fenced for a plant species removal experiment. Dominant or common plant species were removed either individually or in combination several times per month each year from 2014 on. Experimental treatments used in this study were: (1) no plant removal (control); (2) removal of *K. pygmaea* (Kp); (3) removal of *P. saundersiana* L. (Ps); and (4) removal of both (Kp+Ps). Each treatment, applied to 1.5 × 1.5 m plots, was replicated eight times. Plots were randomly distributed and interspaced by approx. 3 m. Urea was applied to half of the plots (146.7 g per plot) once annually in mid-July (Fig. S1), which translates to 300 kg N ha^−1^ yr^−1^ and represents a high fertilization regime (Sun et al. 2008).

### Plant and soil sampling

During the peak of the growing season (August 19^th^–21^st^, 2021), the grid method was used to assess plant diversity in each plot (Jentsch et al. 2012). Here, 1 m × 1 m subplots were analyzed, which were further subdivided into 5 cm × 5 cm squares. Plant species were recorded at the nodes in the upper right corner of each square, resulting in 400 points recorded per plot (Table S1).

Soil attached to roots, i.e., rhizosphere soil, of random plants was sampled by thoroughly shaking the plants and collecting the detached soil on July 1^st^, August 13^th^, and October 26^th^ 2021. From each plot, 4–5 random subsamples were collected down to a depth of 10–20 cm using a 2.5-cm diameter corer. Soils from individual subsamples were thoroughly mixed on site to form one composite sample per plot and transported within 10 min on ice to the laboratory. Upon arrival, each sample was cleaned from larger gravel, stones, plant fragments, and roots and thereafter subdivided into two portions. One portion was sieved through a 2-mm sieve for downstream molecular analyses, the other was air-dried for analysis of soil physicochemical parameters.

Prior to measuring soil properties, soil samples were oven-dried overnight at 45°C. Soil pH was measured using a WTW pH 720 meter (Xylem Analytics Germany Sales GmbH & Co. KG, Weilheim, Germany) in a soil suspension prepared by mixing 5 g of soil with 25 mL of 0.01 M CaCl₂ (Merck KGaA, Darmstadt, Germany). Total organic carbon (TOC) was analyzed using the Vario TOC Cube (Elementar Analysensysteme GmbH, Langenselbold, Germany), and total organic nitrogen (TON) was measured using the APNA-370 nitrogen oxide analyzer (HORIBA, Ltd., Kyoto, Japan). Total ammonium (NH_4_^+^ + NH_3_) and combined nitrate + nitrite concentrations were determined with a Continuous Flow Analyser “Skalar San ++” (Skalar Analytical B.V., Breda, Netherlands).

### DNA extraction and 16S rRNA gene amplicon sequencing

DNA was extracted using the Fast DNA Spin Kit for Soil (Fisher Scientific GmbH, Heiligen, Germany), following the manufacturer’s instructions. DNA concentrations were measured fluorometrically using a Qubit 4 Fluorometer (Thermo Fisher Scientific, Waltham, Massachusetts, USA). 16S rRNA genes of the total bacterial community were amplified using universal bacterial primers 515-f (5’-GTGCCAGCMGCCGCGGTAA-3’) and 907-r (5’-CCGTCAATTCCTTTGAGTTT-3’) (Edwards et al. 2015). Amplification was performed in a 50-µL reaction mixture containing 25 µL of 2-fold Platinum™ Hot Start PCR Master Mix (Fisher Scientific GmbH, Bremen, Germany), 2.5 µL of each forward and reverse primer with adapters for Illumina library preparation (10 µM), 5 µL of DNA template (1 ng µl^_1^), and 15 µL of DNA-free water (PCR grade) (Molzym GmbH and Co. KG, Bremen, Germany). The first amplification step involved an initial denaturation at 94°C for 3 min, followed by 10 cycles of denaturation at 94°C for 45 s, annealing at 50°C for 60 s, and extension at 72°C for 90 s, with a final extension at 72°C for 10 min. PCR products were then purified using Agencourt AMPure XP magnetic beads (Beckman Coulter, Indianapolis, IN, USA). A second PCR was performed to introduce barcodes for Illumina sequencing, using 20 µL of the purified product as the template, with 14 cycles under the same conditions as the first PCR. Equimolar amounts of resulting PCR products were pooled, and amplicon sequencing was carried out using a MiSeq Reagent Kit v3 (Illumina, San Diego, CA, USA).

Raw sequencing reads were processed with QIIME2 version 2023.5 (qiime2.org) (Boylen et al., 2019), including DADA2 for denoising, quality control, chimera removal, and generation of amplicon sequencing variants (ASVs) (Callahan et al. 2016). Taxonomy was assigned using the SILVA database 138.2. (Quast et al. 2012). Processed samples had on average 77,089 paired-end reads, with 90% of all samples ranging between 43,777 and 161,774 paired-end reads (5% and 95% quantiles, respectively). This resulted in a total of 7,317 ASVs across 96 samples. Rarefaction curve analysis indicated that sequencing coverage was sufficient (Fig. S2). This was corroborated by a sequencing coverage of 1 for all samples as assessed using the R package iNEXT v.3.0.0 (Hsieh et al. 2016).

### Quantitative PCR

Quantitative PCR (qPCR) was conducted using the universal primers 1389-f (5′LTGYACACACCGCCCGTL3′) and 1492-r (5′LGGYTACCTTGTTACGACTTL3′) (Tamang et al. 2024). DNA samples were diluted to ca. 1 ng µl^_1^ to standardize template concentrations. The PCR was then performed in a 20-µL reaction mixture containing 10 μL of LightCycler® 480 SYBR Green I Master Mix (Roche, Penzberg, Germany), 0.4 μL of each forward and reverse primer (10 μM), 5 μL of template DNA, and 4.2 μL of DNA-free water (PCR-grade) (Molzym GmbH and Co. KG, Bremen, Germany). An amplified 16S rRNA gene fragment from *Escherichia coli* strain K12, cloned into the pCR 4-TOPO vector (Invitrogen, Waltham, Massachusetts, USA), was used to create a standard curve ranging from 10² to 10L copies µLL¹ (R² = 99.3). Amplification was performed using an initial denaturation step at 95°C for 10 min, followed by 46 cycles consisting of denaturation at 95°C for 30 s, annealing at 52°C for 30 s, and elongation at 72°C for 30 s. A subsequent melting curve analysis was routinely done to check PCR specificity. A dilution series of template DNA confirmed no partial PCR inhibition by co-extracted compounds. All qPCR reactions were conducted in a qTOWER^3^G using the qPCR soft 4.1 software (Analytik Jena GmbH & Co. KG, Jena, Germany). PCR efficiencies were on average 0.84 ± 0.07 (mean ± st. dev.).

### Statistical analysis

Statistical analyses and all visualizations were done in R (R-Core-Team 2025). Alpha diversity was assessed using Hill numbers via the hilldiv R package v.1.5.1 (Alberdi and Gilbert 2019) . Hill numbers provide advantages over traditional diversity indices due to their unified metric that considers both ASV richness and evenness, allowing for a better understanding of community structure (Alberdi and Gilbert 2019). Briefly, ^0^D represents species richness, ^1^D reflects abundant species, and ^2^D highlights dominant species. Evenness was assessed via the alpha-gambin value (McGill et al. 2007) using the R package gambin v. 2.5.0 (Matthews et al. 2014). A higher alpha-gambin value reflects a more even species abundance distribution (McGill et al. 2007).

To evaluate the effects of plant species removal, urea addition, and growing season duration on soil properties, plant species richness, and soil bacterial alpha diversity, we utilized linear mixed-effect models using the *lmer* function of the lme4 package v.1.1-36 (Bates et al. 2015). Results of best fitting model selection are given in Tables S2–S4. Assumptions of the best fitting model were verified by plotting theoretical vs. sample quantiles (Q-Q plot) and residuals versus fitted values using ggplot2 v.3.5.1 (Villanueva and Chen 2019) (Fig. S3-S5). The direction and magnitude of treatment effects was assessed by the effect size β, which is represented by the LMM regression coefficients. Subsequently, the model outcome was tested for significant effects using a Wald type II χ² test using the function Anova within the R package car v.3.1-3. (Fox 2019). Beta diversity changes according to treatments were visualized using nonmetric multidimensional scaling (NMDS) ordination as based on the unweighted UniFrac distance (Lozupone and Knight 2005). Permutational multivariate analysis of variance (PERMANOVA) was conducted to quantify how much of the variance in beta diversity was explained by individual treatment variables and their interactions. This was done using the adonis function in the vegan package v.2.6-10 (Oksanen et al. 2026).

Correlations among individual soil properties and plant richness was tested by Pearson correlation using the vegan package v.2.6-10 (Oksanen et al. 2026). In parallel, the effect of individual soil properties or plant richness on bacterial alpha or beta diversity was tested by linear mixed-effect modeling using the *lmer* function of the lme4 package v.1.1-36 (Bates et al. 2015), with sampling plot as random intercept effect. This was done with and without plant removal, urea addition and their interaction as fixed effects. Model selection is given in Supplementary Table S5 and S6, and the verification of the best fitting model as based on residual and Q-Q plots is shown in Supplementary Fig. S6 and S7. Since Hill numbers ^0^ , ^1^ , and ^2^D were highly correlated with each other (Pearson’s *r* 0.66 - 0.89), ^1^D was used to represent bacterial alpha diversity. For beta diversity, the first dimension of the NMDS analysis was used. Linear mixed modeling analysis was verified by random forest analysis using the rfPermute package v.2.5.2 (Kolisnik et al. 2025).

Subsequently, partial least squares structural equation modeling (PLS-SEM) was performed to discern direct from indirect effects of experimental treatments and identify key soil property predictors of bacterial alpha and beta diversity in soil. This was done using the plspm package v.0.5.1 (Gaston Sanchez 2025). Prior to conducting the PLS-SEM analysis, the normality of the dataset was evaluated by computing the skewness of each variable using the e1071 package v.1.7-16 in R (Dimitriadou et al. 2008). Variables with an absolute skewness value >1 were considered to exhibit substantial non-normality. To address this issue, log transformation was applied to these non-normality variables in order to reduce skewness and approximate a more symmetric distribution (Kock and Hadaya 2018). Although PLS-SEM is a variance-based approach that does not require strict normality assumptions, addressing extreme skewness can improve the reliability of parameter estimates and enhance the model’s predictive accuracy and interpretability. Finally, differential abundance of bacterial ASVs among different treatments was assessed using the DESeq2 package v.1.40.2 (Love et al. 2014).

## Results

### Species loss causes strong shifts in grassland plant diversity

Effects of plant species loss and strong N fertilization on the plant community were assessed after seven continuous years of treatment at the peak of the growing season (August 2021). Both, urea addition and plant species removal had significant effects (*p* < 0.05) on the plant community (Fig. 1). Urea addition significantly decreased plant species richness when compared to control plots (β = –0.500, *p* = 0.002). Plant species removal showed heterogeneous effect sizes (β = –2.250 to 1.500, *p* = 0.047) but overall significantly reduced plant species richness beyond the removed species (Table S7). Here, the strongest decline was observed following the removal of *K. pygmaea* or the combined removal of *K. pygmea* and *P. saundersiana*, whereas a negative effect was absent or less pronounced following removal of *P. saundersiana* alone (Fig. 1a).

**Fig. 1.**
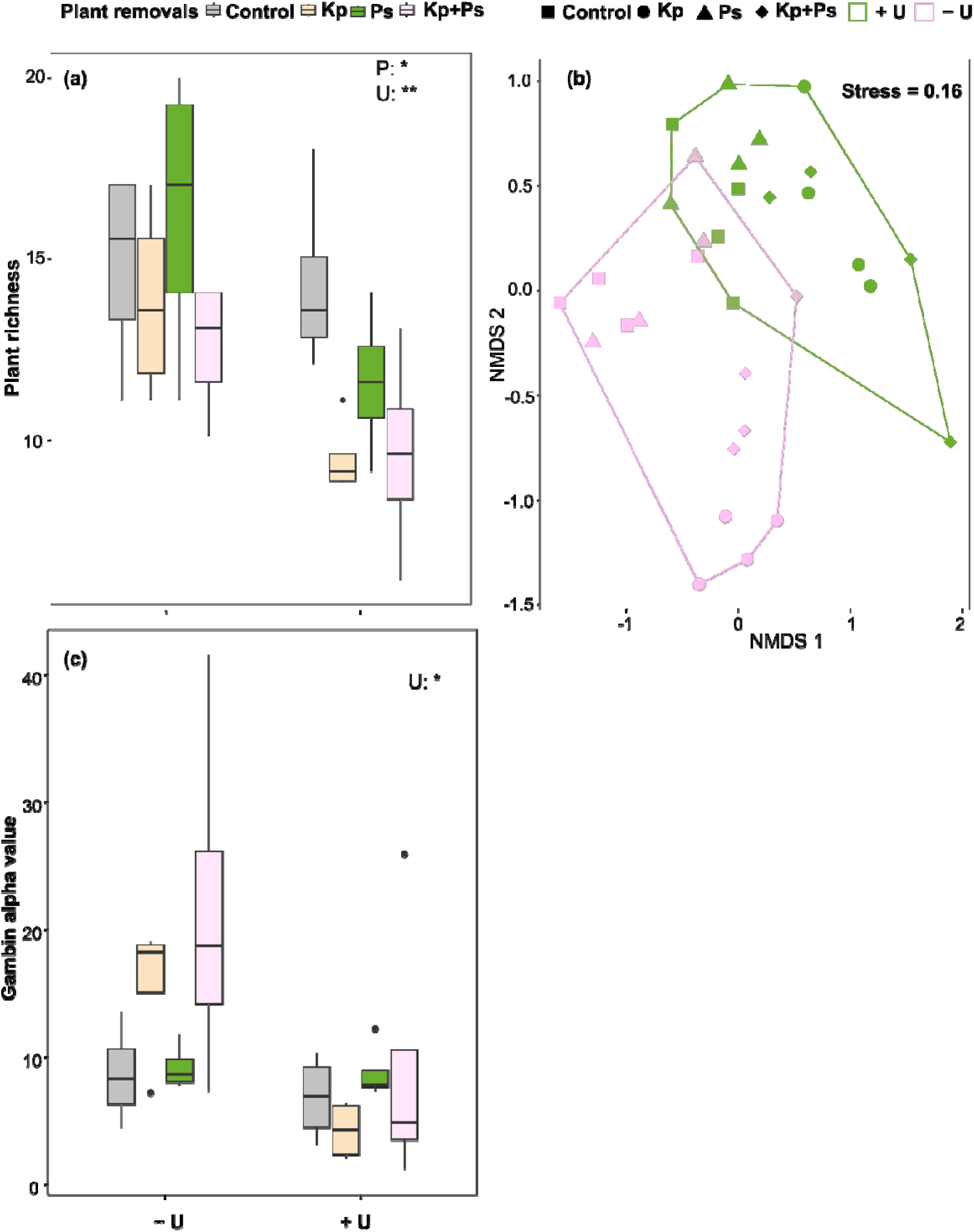
Effects of plant species removal (P) and urea addition (U) on plant communities. (a) Changes in plant species richness according to applied treatments. Treatment effects were tested by linear mixed modelling (lmm), significance values are indicated within the plot: *p* < 0.05 (*), *p* < 0.01 (**), *p* < 0.001 (***). Details of the linear mixed model are given in Supplementary Table S3 and Supplementary Figure S5. (b) Changes in plant beta diversity in respect to applied treatments visualized by non-metric multidimensional scaling (NMDS) ordination as based on Bray-Curtis dissimilarity. (c) Effects of plant species removal and urea addition on plant diversity as indicated by alpha-gambin values. Plant species removal treatments include no removal (control), removal of *Kobresia pygmaea* (Kp), removal of *Potentilla saundersiana* (Ps), and removal of both plant species (Kp+Ps). Urea treatments are indicated as -U (without urea) and +U (with urea).

The composition of the plant community was significantly affected by both treatments as well. Plant communities clearly separated in a NMDS analysis according to treatment, which was pronounced for both urea addition and plant species removal (Fig. 1b). This shift was highly significant as determined by a PERMANOVA analysis (Table 1). Urea addition and plant species removal contributed individually to a similar extent to the variation in plant community composition, explaining 21% and 27% of the variation, respectively. The absence of a significant interaction between urea addition and plant species removal (Table 1) indicated that plant communities did not respond differently to urea addition depending on plant species removal.

**Table 1.** PERMANOVA of the effects of plant species removal, urea addition, and their interaction on plant beta diversity. Df: degrees of freedom; R^2^: proportion of variation explained; F: F-value by permutation, *p*: p-values based on more than 9,000 permutations (the lowest possible *p* is 0.0001).

| Factors | Df | $R^2$ | F | $p$ |
| --- | --- | --- | --- | --- |
| Plant removal | 3 | 0.27335 | 4.8754 | **0.001** *** |
| Urea addition | 1 | 0.20529 | 10.9841 | **0.001** *** |
| Plant removal □ Urea addition | 3 | 0.07282 | 1.2988 | 0.218 |
| Residual | 24 | 0.44854 |  |  |
| Total | 31 | 1 |  |  |

Note: \*, \*\*, \*\*\* indicate significance at $p < 0.05$ , $p < 0.01$ and $p \leq 0.001$ levels, respectively.

The evenness of the plant community, as represented by the alpha-gambin value, changed only significantly under urea addition (β = 6.513, *p* = 0.038, Table S7) and caused a decrease in evenness. Although plant removal caused no apparent significant effect, a clear trend towards a higher evenness of the plant community under *K. pygmaea* removal, or the combined removal of *K. pygmea* and *P. saundersiana* in the absence of urea, was visible (Fig. 1c). This was to be expected considering that *K. pygmea* is the overall dominating plant species in the control plots (Table S1).

### Urea addition and plant species removal result in grassland soil acidification

Effects of urea addition and plant species removal on soil properties were assessed at the start (July), peak (August), and end (October) of the growing season. Soil pH was strongly affected by both, the time of the growing season (β = 0.145 to 0.485, *p* < 0.001) and urea addition (β = –0.400, *p* < 0.001). Specifically, urea addition caused a strong decrease in soil pH by approximately one pH unit during the vegetative period. Plant species removal also had an effect on decreasing soil pH (β = –0.460 to –0.300, *p* = 0.006), albeit less pronounced (Fig. 2, Table S8).

**Fig. 2.**
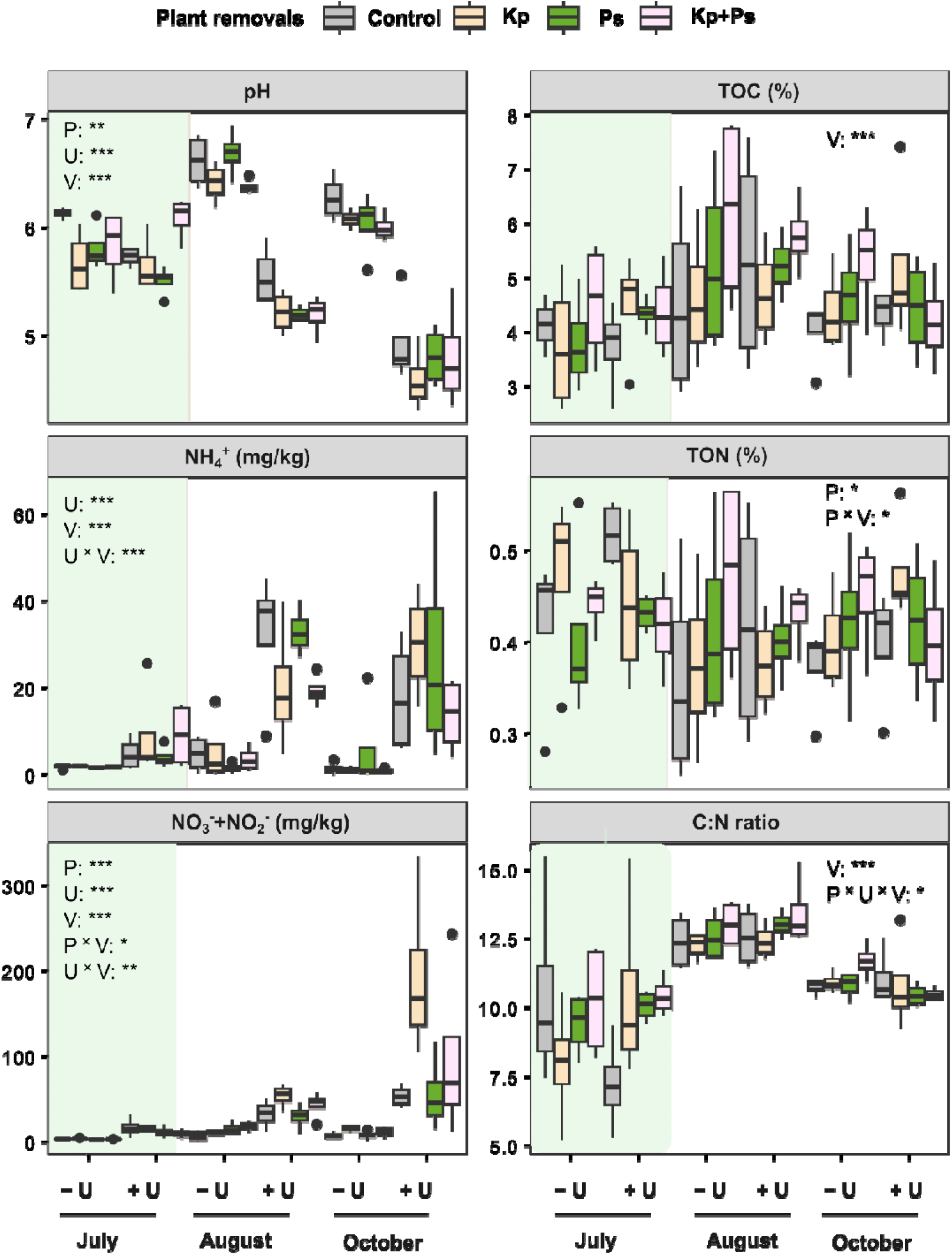
Effect of plant species removal, with and without urea addition, on soil properties across the growing season. The treatments include: no plant removal (Control), removal of *Kobresia pygmaea* (Kp), removal of *Potentilla saundersiana L.* (Ps), and removal of both plants (Kp+Ps). Asterisks denote significant effects of dominant plant removal (P), urea addition (U), and seasonal variation (V) on the plots [\*\*\**p* < 0.001, \*\**p* < 0.01, \**p* < 0.05]. Details are given in Supplementary Table S8. NH_4_^+^ represents ammonium; NO ^-^ + NO ^-^ represent nitrate and nitrite; TOC is total organic carbon; TON is total organic nitrogen; and the C:N ratio refers to the ratio of TOC to TON. Urea treatments are indicated as -U (without urea) and +U (with urea).

Soil NH_4_^+^ and NO₂⁻/NO₃⁻ responded differently across treatments. Soil NH_4_^+^ was positively affected by urea addition (β = 0.554, *p* < 0.001), which was most pronounced in July and decreased in magnitude in October. In contrast, the positive effect (β = 1.232, *p* < 0.001) of urea addition on NO₂⁻/ NO₃⁻ increased throughout the growing season. Here, plant species removal had generally a positive effect as well (β = –0.187 to 0.086, *p* < 0.001). For both, soil NH_4_^+^ and NO₂⁻/NO₃⁻ a significant interaction of urea addition and the growing seasons was observed, while an interaction of plant species removal with the growing season was only significant for NO₂⁻/NO₃⁻ (Fig. 2, Table S8).

In addition, treatment effects were evaluated in respect to TOC, TON, and their ratio (TOC: TON). TOC was significantly affected only by the time of the growing season but with heterogeneous effect sizes (β = –0.109 to 0.406, *p* < 0.001), showing a peak in August throughout all treatment combinations. TON was weakly but significantly affected by the growing season (β = –0.057 to –0.044, *p* = 0.036) and the interaction of plant species removal and growing season (β = –0.039 to 0.089, *p* = 0.037), albeit a clear trend was not discernable. The ratio of TOC:TON was again strongly affected by the time of the growing season (β = 0.308 to 1.939, *p* < 0.001), showing a strong increase in August and a decline thereafter throughout all treatment combinations. However, there was also a weak but significant effect (β = –6.136 to –3.169, *p* = 0.043) of the interaction of plant species removal, urea addition, and time of the growing season (Fig. 2, Table S8).

### Urea addition but not plant species removal negatively affects bacterial alpha diversity

Changes in the abundance and alpha diversity of the bacterial soil community influenced by plant roots (rhizosphere soil) were assessed by 16S rRNA gene qPCR and amplicon sequencing, respectively. Both, the time of the growing season and urea addition had individually a strong and significant effect on total bacterial abundances (Fig. 3). While the duration of the growing season positively affected bacterial abundance (β = 0.030 to 0.032, *p* < 0.001) peaking in August, urea addition had an overall negative effect (β = –0.016, *p* < 0.001). In addition, the time of the growing season showed a weak positive interaction with plant species removal (β = 0.001 to 0.053, *p* = 0.003). A weak significant interaction of growing season, urea addition, and plant species removal was observed as well, albeit with heterogenous effect sizes (β = –0.043 to 0.015, *p* = 0.032) (Fig. 3, Table S9).

**Fig. 3.**
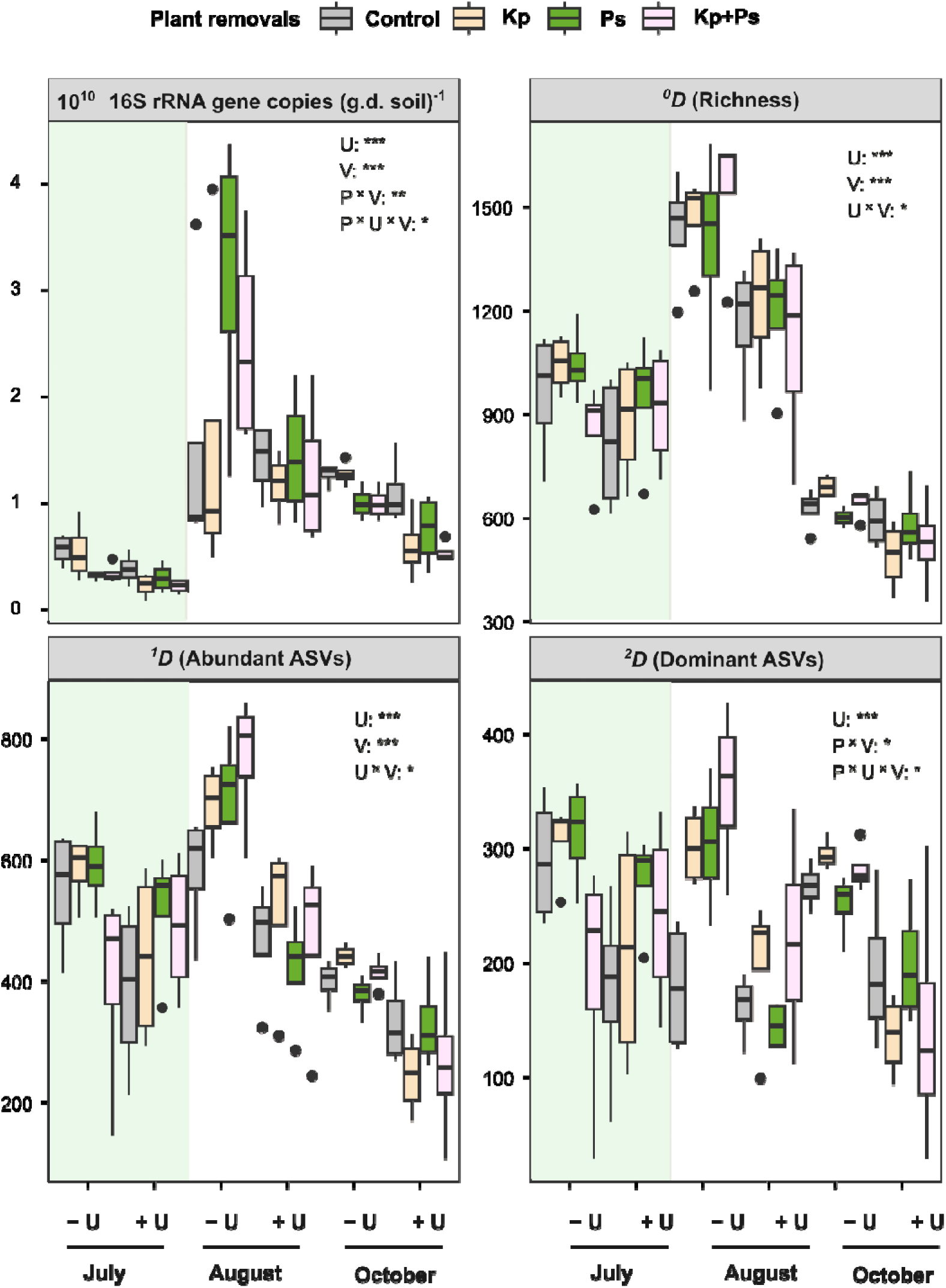
Effects of plant species removal and urea addition on the abundance and alpha diversity of the total bacterial community across the growing seasons. Plant species removal treatments include no plant species removal (Control), removal of *Kobresia pygmaea* (Kp), removal of *Potentilla saundersiana* (Ps), and removal of both species (Kp+Ps). Urea treatments are represented as −U (without urea) and +U (with urea). Asterisks indicate significant effects of plant species removal (P), urea addition (U), and seasonal variation (V) on soil properties (\*\*\**p* < 0.001, ** *p* < 0.01, * *p* < 0.05). Details are given in Supplementary Table S9.

Bacterial ASV richness and abundant ASVs were affected in a similar way by the experimental treatments as total bacterial abundance, albeit without an observed interaction between plant species removal and the growing season (Fig. 3, Table S9). In contrast, dominant ASVs were not affected by the time of the growing season as an individual factor but only by urea addition (β = –113.9, *p* <0.001). Only in interaction with plant species removal (β = –39.9 to 273.5, *p* = 0.012) or in interaction with both plant species removal and urea addition (β = –281.0 to –48.0, *p* = 0.019), a significant but weak effect of the time of the growing season was observed, albeit with heterogenous effect sizes resulting in no clear trend (Fig. 3, Table S9).

### Growing season is the main driver of bacterial community dynamics, with secondary effects of urea addition

Bacterial community dynamics were visualized by a NMDS analysis as based on the unweighted Unifrac distance (Fig. 4a). The latter was chosen to put emphasis on changes in the phylogenetic composition and to detect losses or gains of individual species rather than changes in their relative abundances (Lozupone et al. 2011). Changes in composition were most pronounced in response to the duration of the growing season but also showed a differentiation according to urea addition. These shifts were highly significant as determined by a PERMANOVA analysis (Table 2). The duration of the growing season explained 35% of the variation in the bacterial community composition, while urea addition explained 7%. The interaction of both parameters explained another 7% of the variation, which means that the bacterial community responded differently to urea addition, depending on the month during the growing season (Fig 4a). In particular, the variation explained by urea addition was increasing throughout the vegetative period (Fig. 4b, Table S10). Plant species removal had no significant effect on changes in the bacterial community composition, neither alone, nor in interaction with other parameters (Fig. 4a, Table 2, Table S10).

**Fig. 4.**
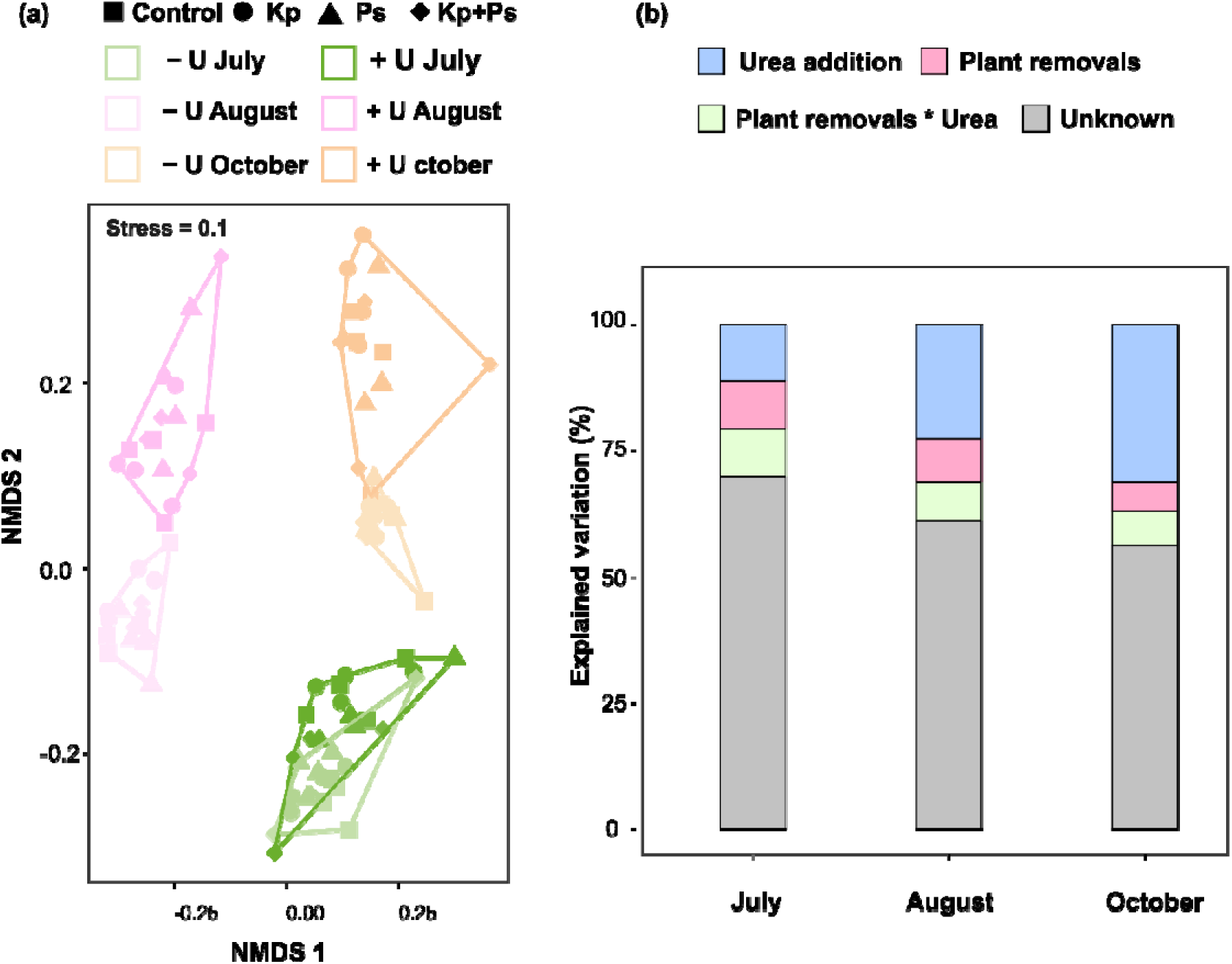
Effects of plant species removal and urea addition on bacterial beta diversity. (a) Nonmetric multidimensional scaling (NMDS) ordination as based on unweighted UniFrac dissimilarity showing changes in bacterial beta diversity with respect to applied treatments throughout the growing season. (b) Explained variance according to treatments and their interaction deduced individually for each analyzed month by PERMANOVA. Details are given in Supplementary Table S10. Plant species removal treatments include no plant species removal (Control), removal of *Kobresia pygmaea* (Kp), removal of *Potentilla saundersiana* (Ps), and removal of both species (Kp+Ps). Urea treatments are represented as -U (without urea) and +U (with urea).

**Table 2.**
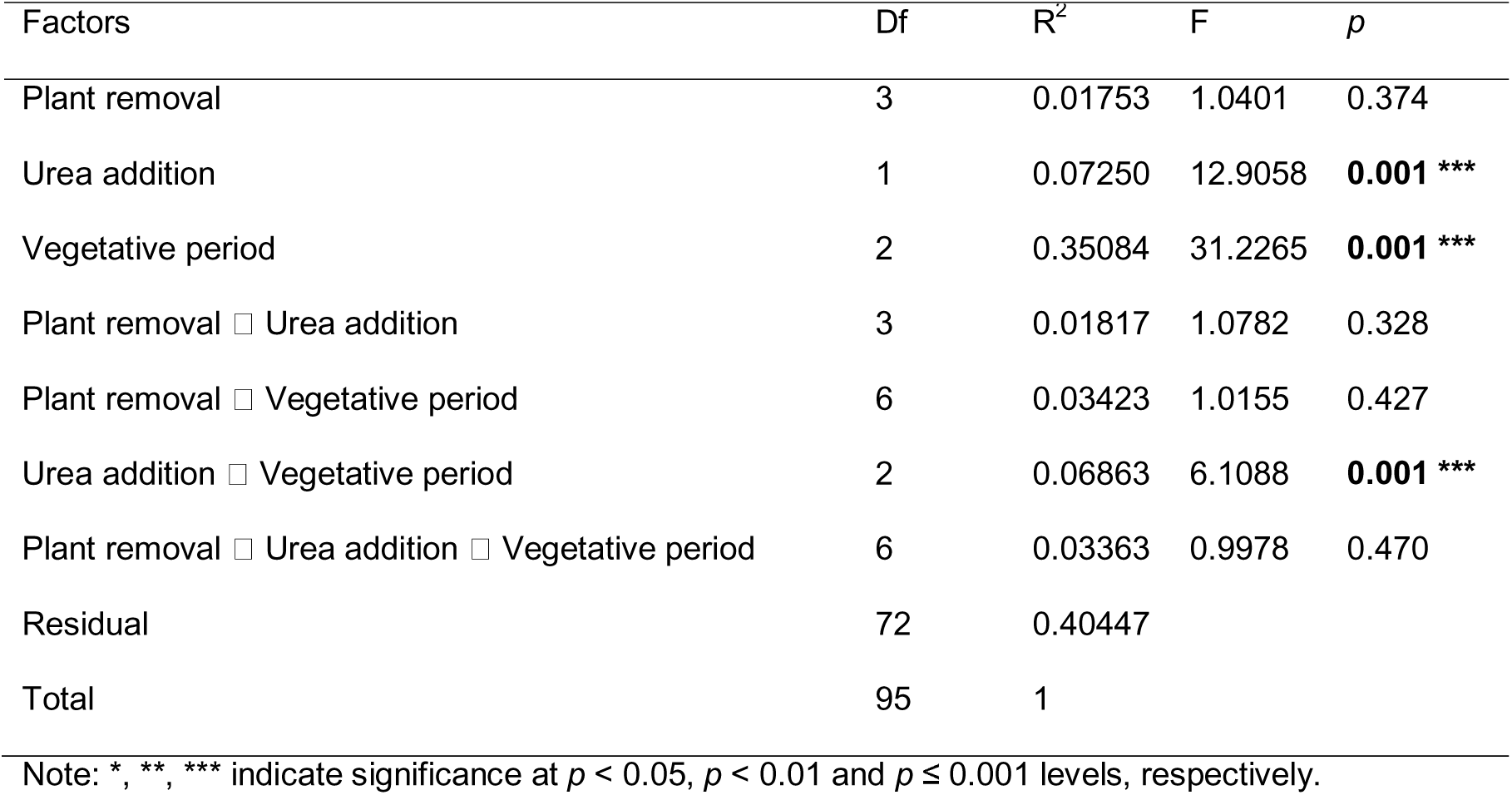
PERMANOVA of the effects of plant species removal, urea addition, the growing season and their interactions on bacterial beta diversity. Df: degrees of freedom; R^2^: proportion of variation explained; F: F-value by permutation, *p*: *p*-values based on more than 9,000 permutations (the lowest possible *p* is 0.0001).

| Factors | Df | $R^2$ | F | $p$ |
| --- | --- | --- | --- | --- |
| Plant removal | 3 | 0.01753 | 1.0401 | 0.374 |
| Urea addition | 1 | 0.07250 | 12.9058 | **0.001** *** |
| Vegetative period | 2 | 0.35084 | 31.2265 | **0.001** *** |
| Plant removal □ Urea addition | 3 | 0.01817 | 1.0782 | 0.328 |
| Plant removal □ Vegetative period | 6 | 0.03423 | 1.0155 | 0.427 |
| Urea addition □ Vegetative period | 2 | 0.06863 | 6.1088 | **0.001** *** |
| Plant removal □ Urea addition □ Vegetative period | 6 | 0.03363 | 0.9978 | 0.470 |
| Residual | 72 | 0.40447 |  |  |
| Total | 95 | 1 |  |  |

Note: \*, \*\*, \*\*\* indicate significance at $p < 0.05$ , $p < 0.01$ and $p \leq 0.001$ levels, respectively.

Rhizosphere soils were dominated by bacteria belonging to the phyla *Actinobacteriota*, *Pseudomonadota*, *Acidobacterota*, *Bacteroidota*, *Bacillota*, *Chloroflexota*, *Planctomycetota*, and *Gemmatimonadota* (in decreasing order, Fig. S8). The relative abundance of the respective phyla was differently affected by the experimental treatments. In the following, their responses are reported in decreasing order of their respective effect size if significant. For instance, *Acidobacteriota*, *Chloroflexota*, *Planctomycetota, and Gemmatimonadota* exhibited significant decreases under urea addition, while *Bacillota*, *Pseudomonadota*, and *Bacteroidota* showed significant increases (Fig. S8, Table S11). Interestingly, these relative abundance changes were not necessarily reflected in the number of significantly enriched and depleted ASVs, which was very balanced across all bacterial phyla (Table S12).

The duration of the growing season had a strong significant effect on the relative abundance of *Pseudomonadota*, *Planctomycetota, Bacillota*, *Chloroflexota* and *Gemmatimonadota*, a weak but significant effect on *Actinobacterota* and no effect on the *Acidobaceriota* and *Bacteroidota* (Fig. S8, Table S11). Significant temporal responses were not necessarily continuous across the vegetative period, but could also show mid-season maxima (e.g., *Gemmatimonadota*) or minima (e.g., *Planctomycetota)* in relative abundances (Fig. S8).

Temporal dynamics across the growing season were also evident at the level of individual ASVs. Between July and August, roughly double as much ASVs were enriched as compared to the ones that were depleted. However, when comparing October to August, roughly three times more ASVs were depleted as compared to the ones enriched (Table S12).

Plant species removal affected only *Gemmatimonadota* directly at the phylum level, showing a positive effect on overall relative abundance (Fig. S8, Table S11). However, if considering the number of significantly enriched or depleted ASVs, plant species removal typically caused a higher number of ASVs to be enriched as compared to those that were depleted. This means that pronounced decreases by a small number of ASVs were counterbalanced by small increases by a larger number of ASVs. At the phylum level, the interaction of plant species removal with the duration of the growing season affected only *Planctomycetota* positively, and its interaction with the duration of the growing season and urea addition negatively affected *Acidobacteriota* and *Planctomycetota* (Fig. S8, Table S11).

### Potential mechanisms differentiating bacterial communities at the peak of the growing season

The peak of the growing season (August) was analyzed in more detail to identify direct and indirect drivers of bacterial diversity. Bacterial alpha diversity was strongly correlated with soil pH, NH₄⁺, and NO₃⁻ + NO₂⁻ (LMMs *r*L=L0.96–0.98, *p*L<L0.001), but not with TON, TOC, TOC:TON ratio, or plant species richness (Fig. 5a). Similarly, bacterial beta diversity was also strongly correlated with soil pH and NO_3_ ^-^ + NO_2_ ^-^ (LMMs rL=L0.97–0.99, pL<L0.001), but not with NH_4_ ^+^, TON, TOC or TOC:TON ratio (Fig. 6a). Random forest analysis confirmed that pH, NH₄⁺, and NO₃⁻+NO₂⁻ were key soil parameters driving bacterial alpha and beta diversity changes (Tables S13 and S14). However, collinearity among these variables was also observed (Fig. 5a and 6a).

**Fig. 5.**
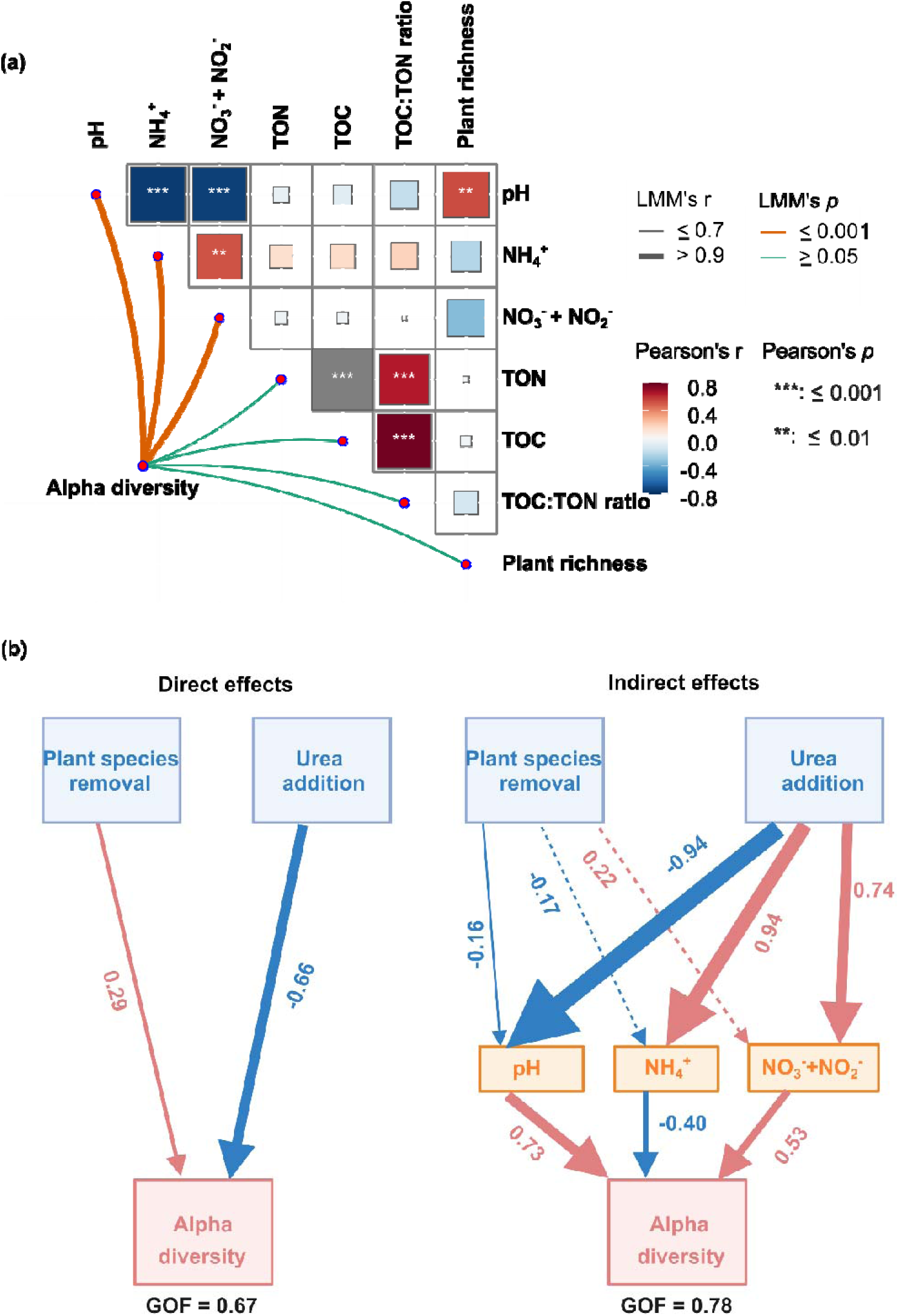
Environmental drivers of bacterial alpha diversity. (a) Correlations between environmental variables and bacterial alpha diversity. Colors represent different correlation types, with solid and dashed lines indicating significant and non-significant correlations, respectively. (b) PLS-SEM illustrating the relationships among experimental treatments, soil variables, and bacterial alpha diversity. Red and blue arrows denote positive and negative relationships, respectively, while solid and dashed lines indicating significant (*p* < 0.05) and non-significant associations. Numbers adjacent to pathway arrows represent standardized path coefficients.

**Fig. 6.**
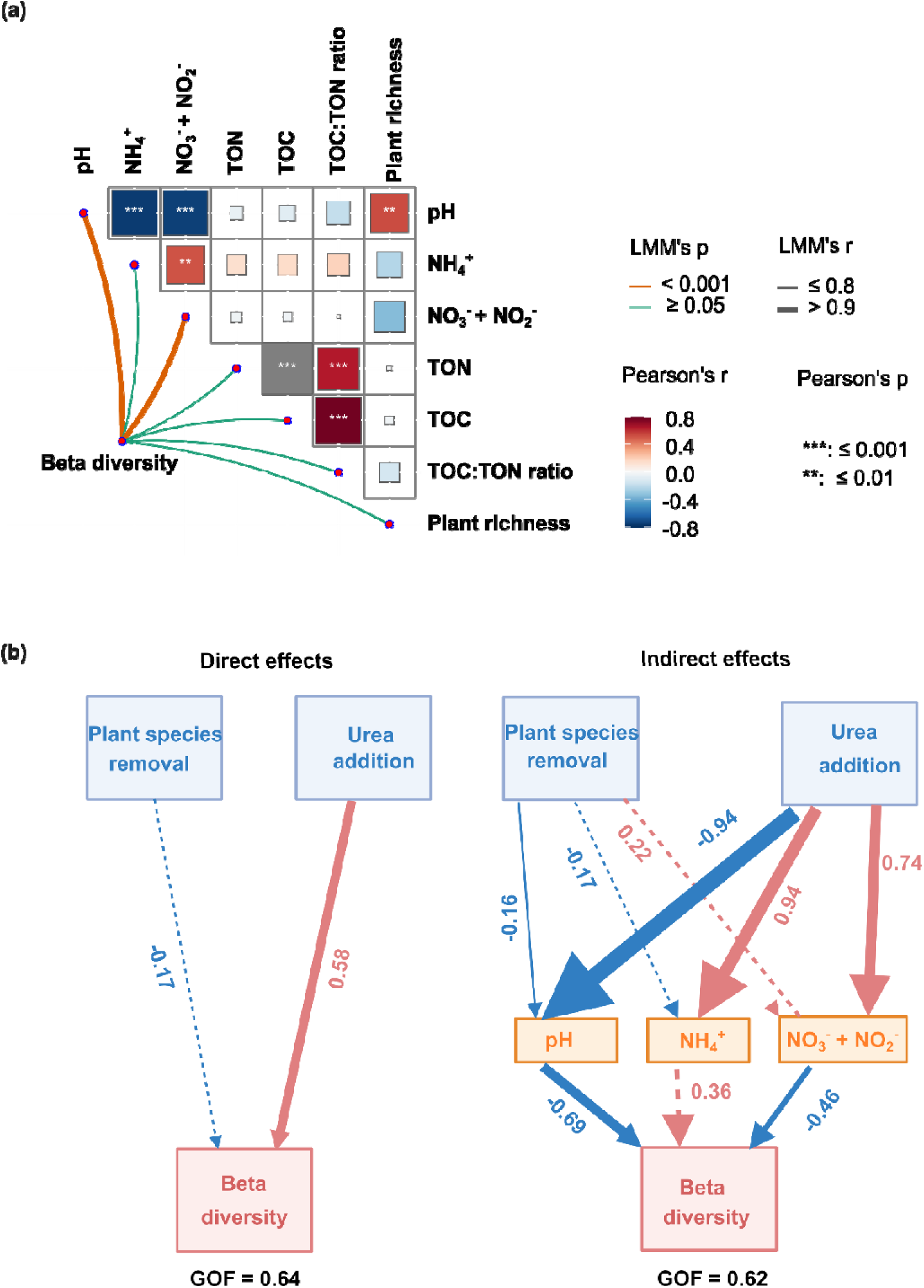
Environmental drivers of bacterial beta diversity. (a) Correlations between environmental variables and bacterial beta diversity. Colors represent different correlation types, with solid and dashed lines indicating significant and non-significant correlations, respectively. (b) PLS-SEM illustrating the relationships among experimental treatments, soil variables, and bacterial beta diversity. Red and blue arrows denote positive and negative relationships, respectively, while solid and dashed lines indicating significant (*p* < 0.05) and non-significant associations (*p* ≥ 0.05). Numbers adjacent to pathway arrows represent standardized path coefficients.

To further disentangle direct from indirect effects of the experimental treatments on bacterial alpha diversity, PLS-SEM analyses were conducted. This included key soil parameters identified by the above correlation analysis. PLS-SEM was chosen because it can deal with small sample sizes (Kock and Hadaya 2018). Our results revealed that direct effects of urea addition and plant species removal on bacterial alpha diversity were both significant (*p* <0.05), with urea addition showing a stronger impact than plant species removal (Fig. 5b). The predominance of urea as a main driver of alpha diversity was consistent with the findings from the linear mixed model analysis (Fig. 3, Table S9). When considering indirect effects via identified key soil parameters, soil pH showed the strongest significant impact on bacterial alpha diversity followed by NO₃⁻ + NO₂⁻ and NH_4_^+^ (Fig. 5b). Soil pH positively affected bacterial alpha diversity but was significantly reduced by urea addition and to a minor extent by plant species removal. NO₃⁻ + NO₂⁻ positively affected bacterial alpha diversity as well, while being positively affected by urea addition only. In contrast, NH_4_^+^ negatively affected bacterial alpha diversity, but was again positively affected by urea addition itself.

Bacterial beta diversity was significantly and positively affected by urea addition, while plant species removal showed no noticeable impact (Fig. 6b). This result is line with the findings from the NMDS and PERMANOVA analyses (Fig. 3a and Table 1). When considering indirect effects through key soil parameters, pH emerged as the primary factor shaping bacterial beta diversity, followed by NO₃⁻ + NO₂⁻. Soil pH exhibited a negative impact on bacterial beta diversity and was significantly reduced by both plant species removal and urea addition. Similarly, NO₃⁻ + NO₂⁻ also had a negative effect on bacterial beta diversity; however, its levels were positively influenced by urea addition.

## Discussion

### Grassland flora biodiversity declines substantially under *Kobresia pygmaea* removal and strong N perturbation

Understanding how co-occurring plant species loss and intense N fertilization affect plant communities and their associated bacterial soil microbiome is crucial for evaluating the impacts of anthropogenic global change drivers on ecosystem functionality. This is important because these elements are essential for supporting nutrient cycling, soil health, and the structural integrity of terrestrial biomes (Bardgett and Van Der Putten 2014, Pugnaire et al. 2019, Cheng et al. 2025). We observed that removing *K. pygmaea* as the dominant plant species in *Kobresia*-dominated grasslands on the Tibetan Plateau led to a significant reduction in overall plant richness (Fig. 1a). This is likely due to its role in forming dense mats that stabilize the soil and retain moisture, creating a favorable microenvironment for other species (Miehe et al. 2008, Miehe et al. 2019). In addition, these mats not only prevent soil erosion but also help retain soil nutrients (Miehe et al. 2008, Zhang et al. 2017, Miehe et al. 2019), which likely supports the growth of other non-dominant plant species as well. This pronounced negative impact on plant species richness was also observed when removing the dominant *K. pygmaea* in combination with the common *Rosaceaea* member *P. saundersiana,* but not if non-dominant *P. saundersiana* was removed alone (Fig. 1a). This emphasizes even more so the ecological importance of *K. pygmaea* for the whole grassland ecosystem. Our results are in line with biodiversity-ecosystem function experiments, which highlighted that in natural systems loss of low-abundance or low-function species has smaller or negligible effects as compared to loss of dominant, high-function species (Smith and Knapp 2003, Gaston 2011, Avolio et al. 2019, Genung et al. 2020).

Increased N input in grasslands has been identified as a significant threat to its biodiversity (Zhang et al. 2021, Wang et al. 2023, Hu et al. 2024), including a decline in plant diversity (Clark and Tilman 2008, Zhang et al. 2014, Band et al. 2022, Zhang et al. 2022). Our study aligns with these findings, revealing that the common N fertilizer urea applied at a high fertilization rate (Sun et al. 2008) significantly reduced plant species richness in the studied alpine grassland (Fig. 1a), accompanied by a significant shift in plant community composition (Fig. 1b). Increasing N input can negatively impact plant diversity in various ways. Abiotic effects include soil acidification (Band et al. 2022), as observed in our study as well (Fig. 2), and toxicity to higher ammonia levels (Britto and Kronzucker 2002). Biotic effects can be attributed to nutrient limitation supporting species coexistence (Harpole and Tilman 2007, Soons et al. 2017). Here, reduction in plant diversity could be attributed to the nutrient imbalance caused by excessive nitrogen, favoring plant species such as grasses adapted to high N availability, while suppressing less competitive species, including forbs and other plants, that rely on lower N availability (Bobbink et al. 2010). Interestingly, we observed no interaction between the strong effect of urea addition and the less strong effect of plant species removal on plant species richness declines (Fig. 1a), indicating that both act via independent mechanisms on plant biodiversity as outlined above.

Our study further showed that seven years of plant species removal had no impact on soil TOC, TON, or ammonium (NH₄⁺) levels, while soil pH as well as combined nitrate and nitrite (NO₃⁻ + NO₂⁻) concentrations were significantly altered. Similar findings have been reported from previous research, which observed that removing plant species from grasslands often has minimal impact on relatively conservative soil properties such as soil C and N pools (Wardle et al. 1999, Marshall et al. 2011, Li et al. 2018, Yang et al. 2021). Interestingly, soil pH significantly decreased with plant removal, which was most pronounced if dominant *K. pygmaea* was removed alone or in combination with *P. saundersiana* (Fig. 2). This is consistent with previous findings, where pH declines were observed after understory vegetation removal in forests (Lei et al. 2021, Zhang et al. 2022), or when studying the effects of individual grassland plant species (Bardgett et al. 1999). However, pH effects can be plant species-specific, with no measurable effects being observed in other grassland plant removal studies (Wardle et al. 1999, Chen et al. 2016). In our study, pH effects may be due to *K. pygmaea*’s contributions to soil organic matter buildup and efficient N uptake (Miehe et al. 2019). With reduced organic input, the soil’s ability to neutralize acidity diminishes, resulting in a lower pH (Amelung W 2018). We also observed that NO₃⁻ + NO₂⁻ concentration increased over time, particularly in October and under *K. pygmaea* removal (Fig. 2). This may be due to the species’ shifting preference from NO₃⁻ towards NH₄⁺ over the vegetative season (Jiang et al. 2016). Our results highlight the critical role of *K. pygmaea* in regulating soil pH and N cycling on alpine meadows on the Tibetan Plateau.

Urea addition strongly impacted soil pH, NH₄⁺, and NO₃⁻ + NO₂⁻ concentrations, even more than plant species removal (Fig. 2). Soil acidification by urea addition confirmed the manifold observations by previous studies (Barak et al. 1997, Fujii et al. 2008, Zhou et al. 2014, Tian and Niu 2015) including Tibetan grasslands (Hu et al. 2024). The increasing negative offset in soil pH over the growing season under urea addition was likely caused by nitrification and subsequent nitrate leaching. The metabolic process of nitrification releases protons (Ward et al. 2011), and when produced NO_3_ ^−^ is leached, base cations are removed concomitantly in neutral and moderately acidic soils (Breemen et al. 1984, De Vries and Breeuwsma 1987). Our results showed that NH₄⁺ concentrations were highest in July, followed by a decline through October, while NO₃⁻ + NO₂⁻ concentrations exhibited an increasing trend over the same period (Fig. 2). This pattern likely reflects the activity of nitrifying microorganisms and is in line with ^15^N-tracer studies following the temporal dynamics of nitrification-driven soil acidification caused by urea addition (Dong et al. 2022).

### Direct and indirect drivers of bacterial belowground responses to plant species loss and strong N perturbation

The soil bacterial community responded mainly to the duration of the growing season and urea addition, while plant removal had a minor effect. This was the case at the level of bacterial abundances, alpha and beta diversity (Fig. 3, Fig. 4). Especially in August, at the peak of the growing season, observed differences were most pronounced. We analyzed this time point in more detail for direct and indirect drivers of plant-soil feedbacks affecting bacterial belowground biodiversity.

Indirect effects via soil acidification were clearly the most dominant factor in decreasing bacterial alpha diversity and driving changes in bacterial beta diversity towards diverging communities (Fig. 5, 6). Decreases in soil pH were driven largely by urea addition and to a minor extend by plant species removal (Fig. 5, 6) as outlined in detail above. Soil pH has been identified before as one of the major drivers of belowground bacterial diversity (Fierer and Jackson 2006, Lauber et al. 2009, Rousk et al. 2010). Most bacterial species typically regulate their intracellular pH to remain within one unit of neutrality (Madigan et al. 2000) and display in pure culture narrow pH ranges permissive for growth, usually varying across 3-4 pH units between minimum and maximum (Rousk et al. 2010). In addition, natural bacterial communities of soils have been shown to be adapted to their *in situ* pH, displaying a 50% reduction in growth at deviations by 1.5 pH units (Bååth 1996, Fernandez-Calvino and Bååth 2010). These narrow pH optima were proposed before to explain the strong selecting pressure of pH on bacterial community composition in soil (Rousk et al. 2010). Changing NO₃⁻ + NO₂⁻ concentrations were the second most important indirect drivers of bacterial alpha and beta diversity. As outlined above, they were increasing solely due to urea addition but not in response to plant species removal (Fig. 5, 6). Increasing NO₃⁻ + NO₂⁻ concentrations caused an increase in bacterial alpha diversity as well as a clear change in bacterial beta diversity towards more similar communities. This is in contrast to a N deposition experiment in the grasslands of Inner Mongolia, China, where effects of N fertilization on bacterial alpha diversity were not observed (Zeng et al. 2016). However, the same study reported a decrease of bacterial alpha diversity in response to increased NH₄⁺ levels. This is in line with our observations, where NH₄⁺ was identified as the third indirect driver negatively affecting bacterial alpha diversity, while having no effect on beta diversity (Fig. 5, 6). Other studies have reported a generally negative effect of both ammonia- and nitrate-based fertilizers on bacterial alpha diversity, while bacterial beta diversity separated in response to the different N sources (Castellano-Hinojosa et al. 2021, Zhang et al. 2021). This could be potentially explained by the stimulation of different microbial guilds such as nitrifiers in the case of NH₄⁺ and denitrifiers in the case of NO₃⁻ (Castellano-Hinojosa et al. 2020), or the differential response of copiotrophs vs. oligotrophs (Fierer et al. 2007) towards different N sources. However, effects of N input, including different N sources on microbial diversity metrics and activity, can vary across sites (Ramirez et al. 2010, Ramirez et al. 2012, Geisseler et al. 2016, Tian et al. 2017, Zhou et al. 2017, Wang et al. 2018) and are difficult to be generalized.

Direct effects of urea addition and plant species removal on bacterial diversity metrics were observed as well (Fig. 5, 6). Here, urea had a strong negative effect on bacterial alpha diversity while increasing bacterial beta diversity. This was in line with previous studies (Zeng et al. 2016, Lu et al. 2021, Wang et al. 2023). Direct effects could be due to increased osmotic potential and ion toxicity (Eno et al. 1955, Omar and Ismail 1999) and the generally higher N availability. Since higher N availability increases plant productivity (LeBauer and Treseder 2008) and production of root exudates (Zhu et al. 2016), fast-responding copiotrophs may benefit over nutrient limitation-adapted oligotrophs within the bacterial soil community (Fierer et al. 2007). This is supported by our observation that bacterial phyla harboring typical oligotrophs such as *Acidobacteriota*, *Chloroflexota*, *Planctomycetota*, and *Gemmatimonadota* (Fierer et al. 2007, Fierer et al. 2012) exhibited significant decreases under urea addition, while phyla harboring typical copiotrophs such as *Bacillota*, *Bacteroidota*, and *Pseudomonadota* (Fierer et al. 2007, Fierer et al. 2012, Zeng et al. 2016) showed notable significant increases (Fig. S8, Table S11). In addition, root exudates can function as signaling compounds in plant-microbe interactions (Coskun et al. 2017), and a changed root exudate profile may directly modulate the soil microbiome (Sasse et al. 2018). With respect to N-cycling microorganisms, N_2_-fixing microorganisms may decline under generally higher N availability (Berthrong et al. 2014, Liao et al. 2021), while other N-cycling guilds such as nitrifiers and denitrifiers would benefit (Hartmann et al. 2013, Castellano-Hinojosa et al. 2020). In addition, differential preferences for urea versus NH₄⁺ have been shown among ammonia oxidizing bacteria and archaea (Qin et al. 2024) and may partially explain direct effects of urea on diversity metrics as well.

Plant species removal had a minor direct effect on bacterial diversity metrics (Fig. 5, 6). Typically, a higher plant diversity is beneficial for microbial belowground diversity (Eisenhauer 2016) and microbial ecosystem functions, including microbial biomass and respiration (Eisenhauer et al. 2010, Chen et al. 2019), as well as soil carbon storage (Lange et al. 2015). In our study, plant species removal rather led to increases in bacterial alpha diversity, while bacterial beta diversity was not directly affected (Fig. 5, 6). Removing plant species may alter the availability of resources and ecological niches in soil (Wardle et al. 1999), and plants are known to modulate their surrounding soil microbiome (Sasse et al. 2018). Removal of the numerically most dominant plant species *K. pygmaea* with and without the common plant species *P. saundersiana* from *Kobresia*-dominated pastures resulted in a trend towards a more even distribution of plant species (Fig. 1c). In turn, this may have opened a more evenly distributed higher niche space for soil microorganisms. This interpretation is supported by our observation that plant species removal resulted in a higher number of phylum-specific ASVs to be enriched as compared to those that were depleted (Fig. S8, Table S12).

## Conclusions

Our study emphasizes the complex impacts of dominant plant species loss and increasing N inputs on ecosystem functions. While the removal of dominant plant species did not directly alter the overall structure of soil bacterial communities, it L along with urea addition L had pronounced effects on the plant community itself as well as on soil properties, such as pH and N availability. These changes subsequently influenced bacterial community dynamics, indicating that soil bacterial responses on the Tibetan Plateau are driven more by shifts via soil chemical parameters than by the direct alterations in plant composition. Our findings suggest that ongoing changes in plant dominance and N input could alter soil nutrient cycling and bacterial functional composition, leading to shifts in belowground bacterial biodiversity towards copiotrophs that may enhance soil carbon turnover and contribute to positive feedbacks on global climate change.

## ACKNOWLEDGEMENTS

This research was conducted within the framework of the DFG-funded Sino-German Research Training Group “Geo-ecosystems in transition on the Tibetan Plateau” (TransTiP, GRK 2309, DFG grant 317513741). We thank Johannes Sikorski for fruitful discussions on R analyses. All sampling and downstream research, including metabarcoding of genetic resources for taxonomic purposes in Germany, was conducted according to the access and benefit-sharing agreement within the corresponding Memorandum of Understanding (signed November 30^th^, 2019) and its addendum (signed September 13^th^, 2021) between the Institute of Tibetan Plateau Research, Chinese Academy of Sciences, and the Technische Universität Braunschweig, Germany.

## Data availability statement

All amplicon sequences were deposited at the Sequence Read Archive at the National Center for Biotechnology Information (NCBI) under the Bioproject number PRJNA1294614. R code is available in GitHub under the following address: https://github.com/wudaidaidsmz/Tibetan-Plateau.

## Conflict of interest statement

All authors confirm that they have no conflict of interest to declare.

## Supplementary Material

### Supplementary Figures

**Fig. S1.**
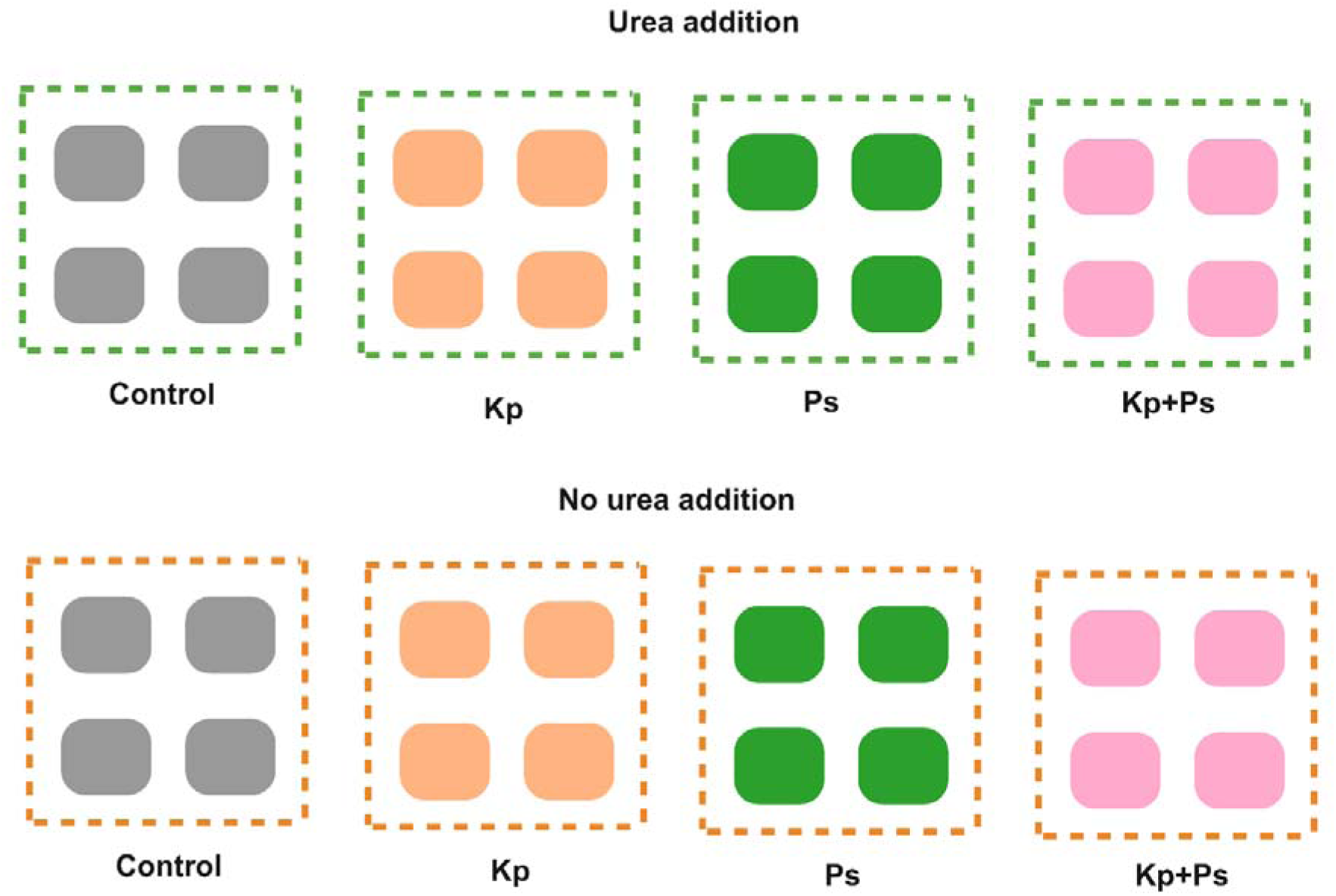
Experimental plots for this study. The treatments include: no plant removal (Control), removal of *Kobresia pygmaea* (Kp), removal of *Potentilla saundersiana L*. (Ps), and removal of both plants (Kp+Ps). Urea addition and no urea addition.

**Fig. S2.**
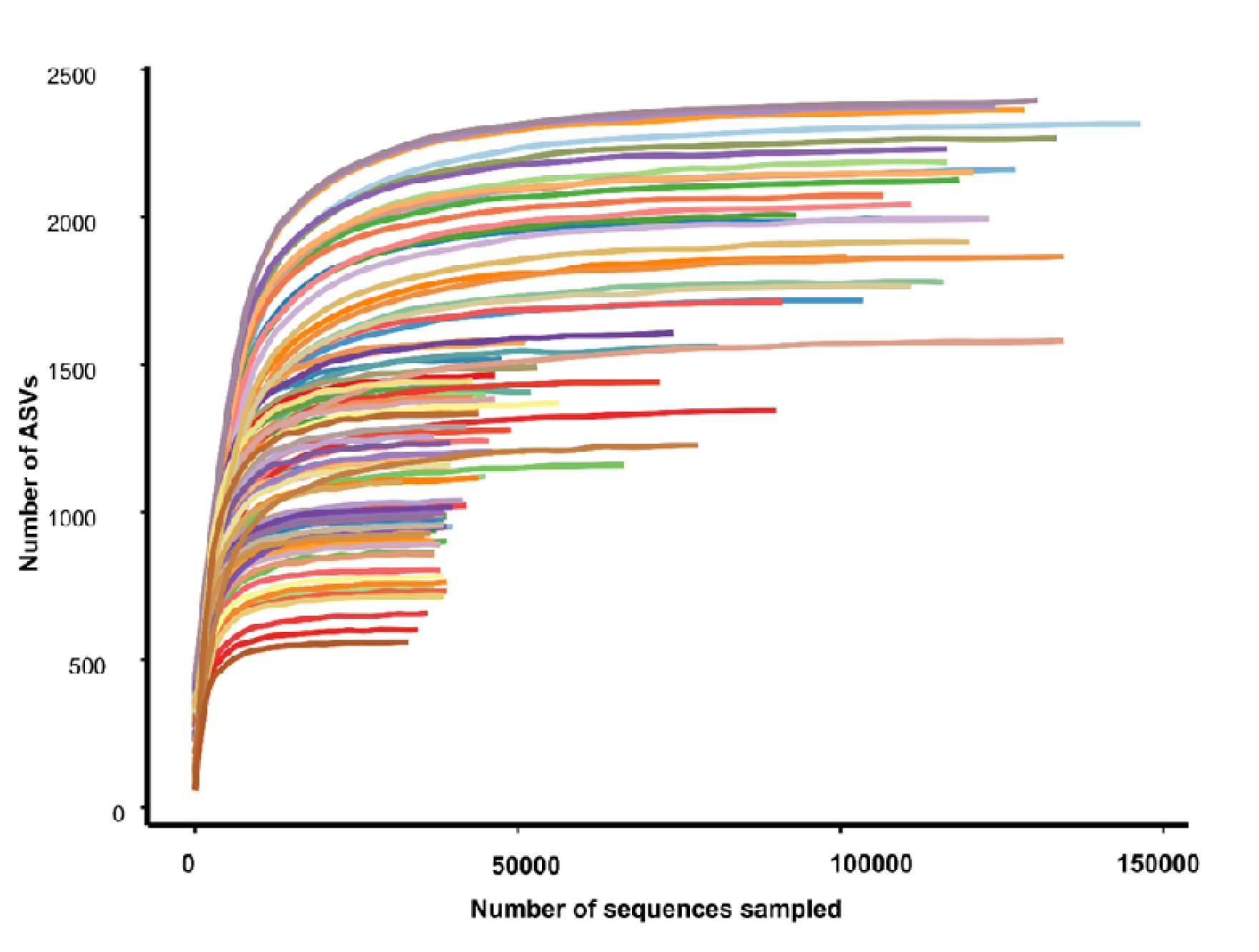
Rarefaction curve analysis of amplicon sequencing variants (ASVs) detected by bacterial 16S rRNA gene amplicon sequencing across all analyzed samples.

**Fig. S3.**
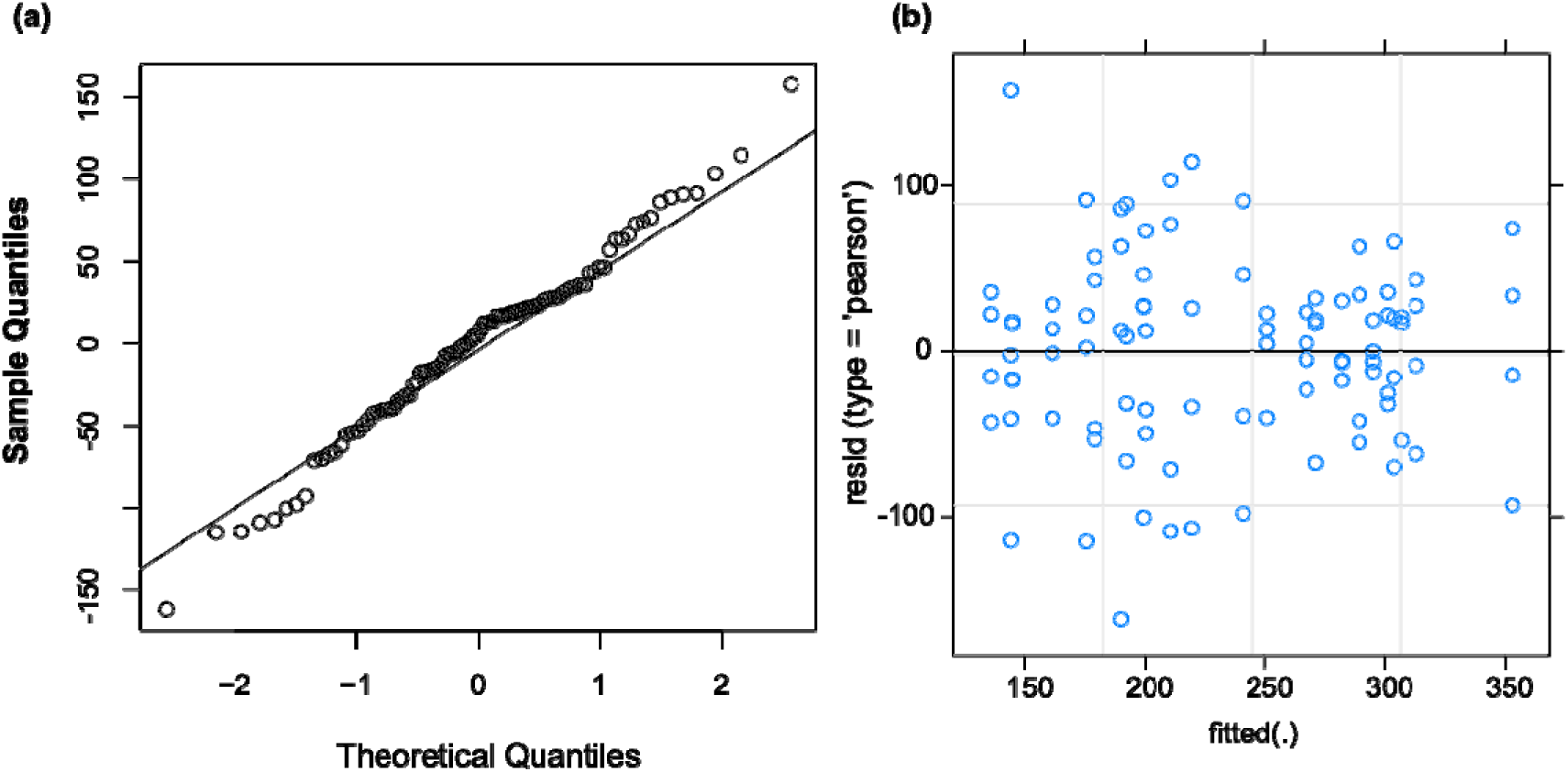
Diagnostic plots for the best-selected Linear Mixed Model (LMM) examining the relationship between bacterial alpha diversity and experimental treatments (plant species removal and urea addition). (a) QQ-Plot of residuals: Assesses the normality of residuals. If the points align closely with the reference line, the residuals are approximately normally distributed. (b) Residuals vs. Fitted Values: Evaluates homoscedasticity. A random scatter indicates constant variance, while visible patterns may suggest heteroscedasticity or model misspecification.

**Fig. S4.**
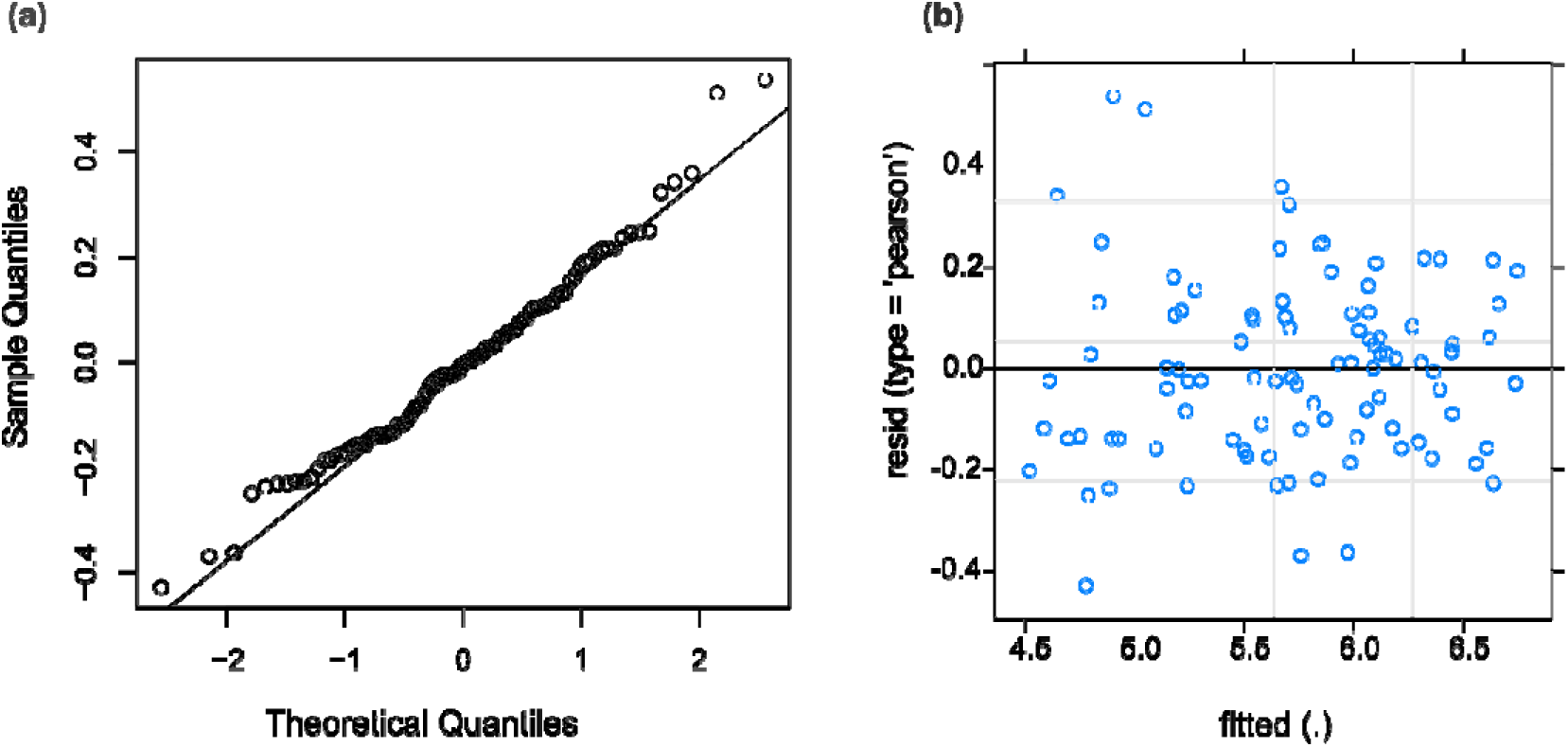
Diagnostic plots for the best-selected Linear Mixed Model (LMM) examining the relationship between soil properties and experimental treatments (plant species removal and urea addition). (a) QQ-Plot of residuals: Assesses the normality of residuals. If the points align closely with the reference line, the residuals are approximately normally distributed. (b) Residuals vs. Fitted Values: Evaluates homoscedasticity. A random scatter indicates constant variance, while visible patterns may suggest heteroscedasticity or model misspecification.

**Fig. S5.**
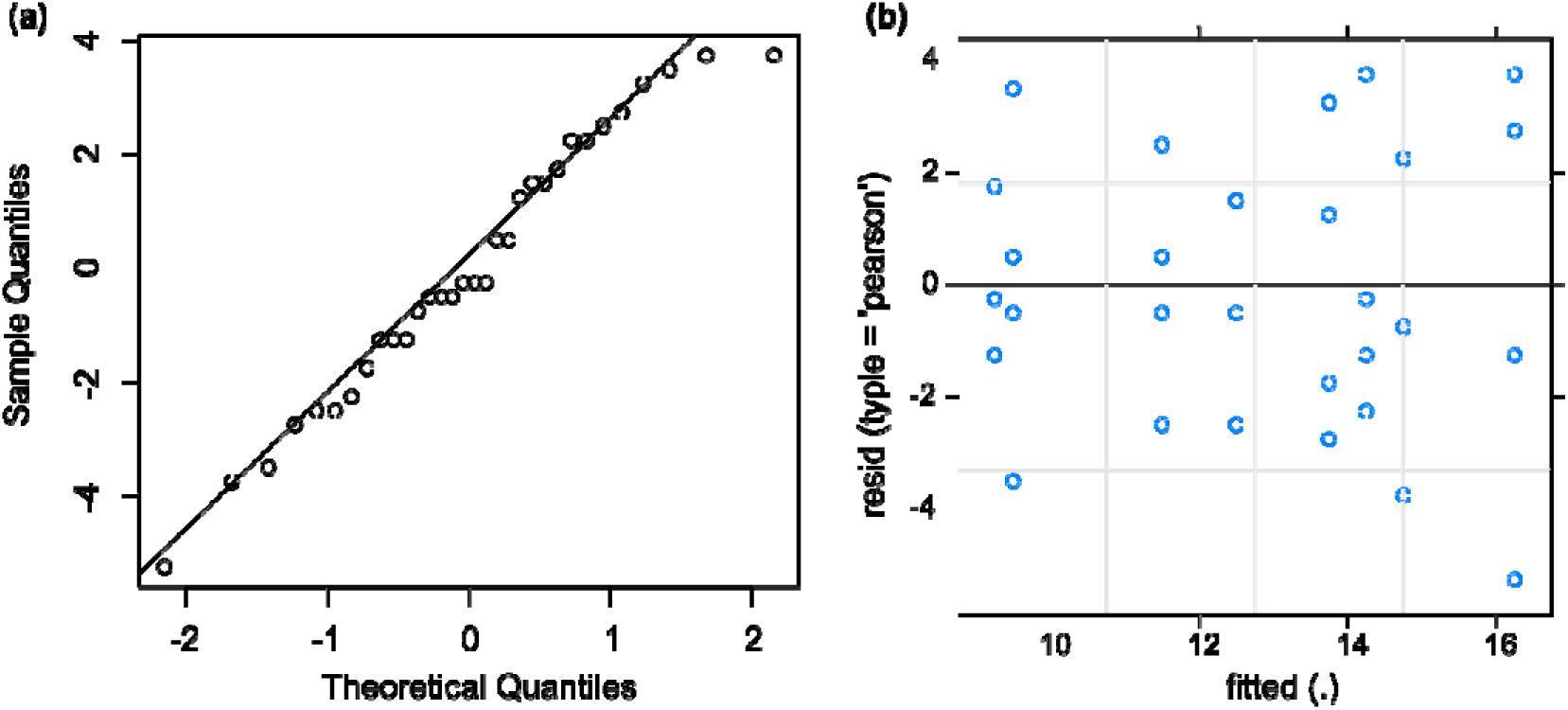
Diagnostic plots for the best-selected Linear Mixed Model (LMM) examining the relationship between plant species richness and experimental treatments (plant species removal and urea addition). (a) QQ-Plot of residuals: Assesses the normality of residuals. If the points align closely with the reference line, the residuals are approximately normally distributed. (b) Residuals vs. Fitted Values: Evaluates homoscedasticity. A random scatter indicates constant variance, while visible patterns may suggest heteroscedasticity or model misspecification.

**Fig. S6.**
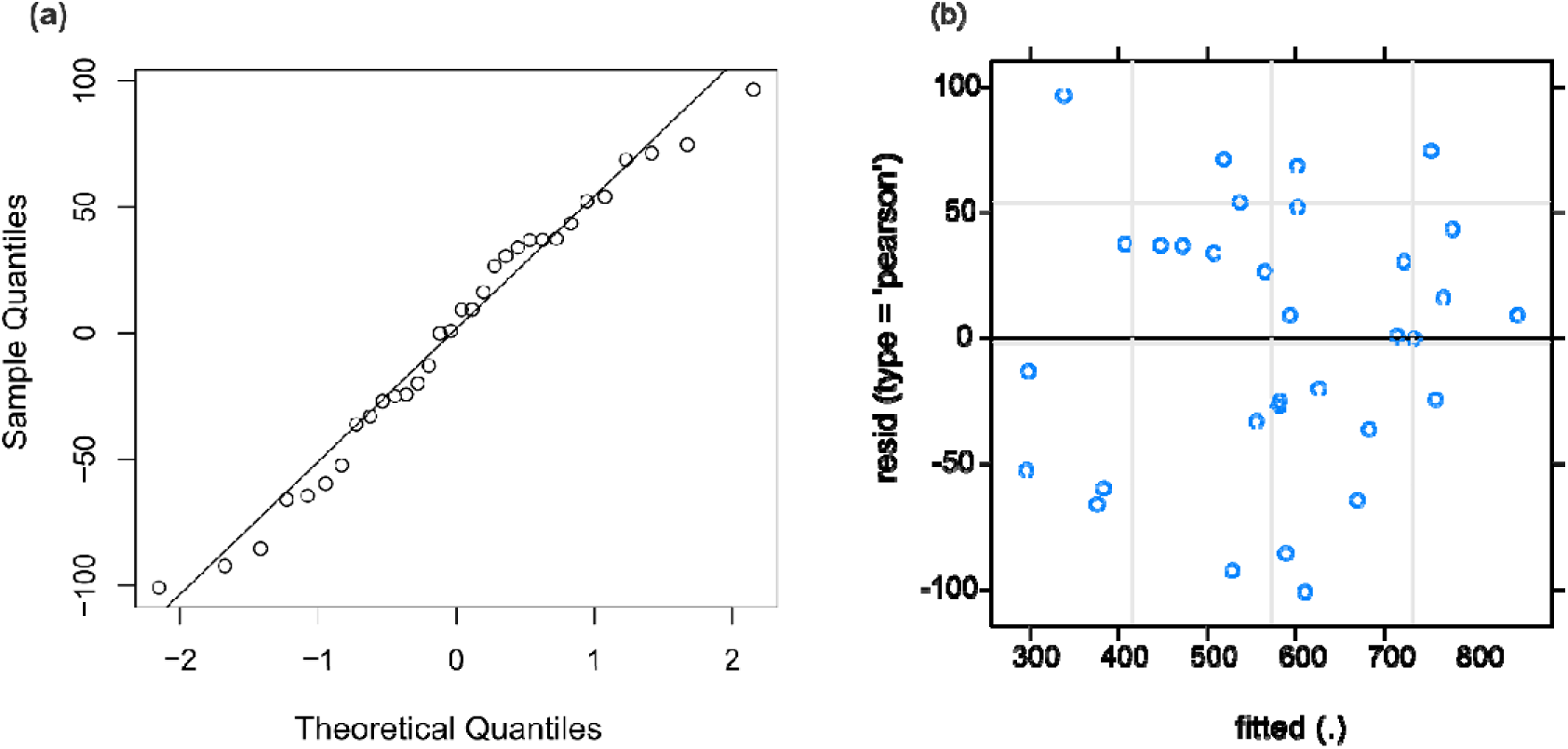
Diagnostic plots for the best-selected Linear Mixed Model (LMM) examining the relationship between soil properties, experimental treatments (plant species removal and urea addition) and bacterial alpha diversity. (a) QQ-Plot of residuals: Assesses the normality of residuals. If the points align closely with the reference line, the residuals are approximately normally distributed. (b) Residuals vs. Fitted Values: Evaluates homoscedasticity. A random scatter indicates constant variance, while visible patterns may suggest heteroscedasticity or model misspecification.

**Fig. S7.**
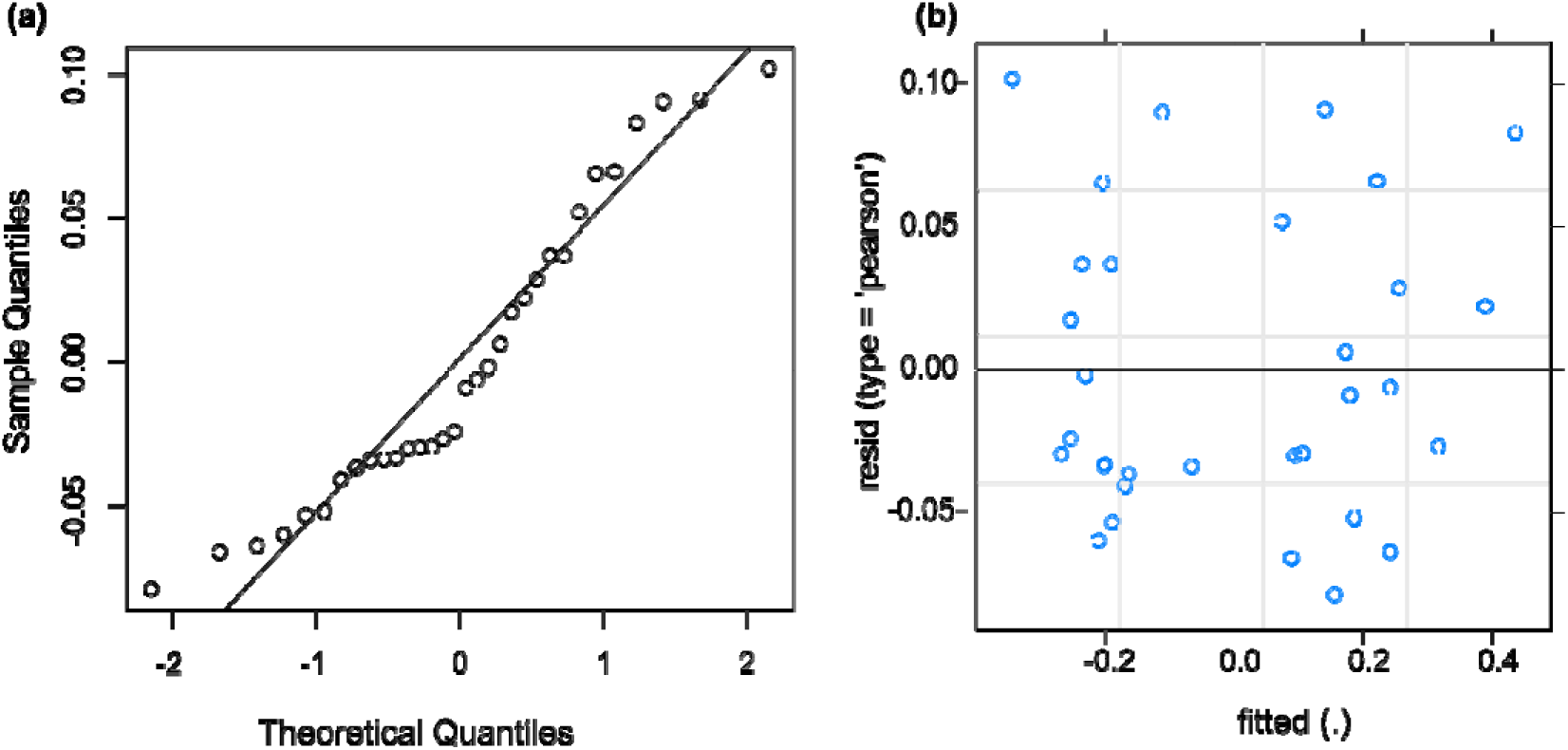
Diagnostic plots for the best-selected Linear Mixed Model (LMM) examining the relationship between soil properties, experimental treatments (plant species removal and urea addition) and bacterial beta diversity. (a) QQ Plot of Residuals: Assesses the normality of residuals. If the points align closely with the reference line, the residuals are approximately normally distributed. (b) Residuals vs. Fitted Values: Evaluates homoscedasticity. A random scatter indicates constant variance, while visible patterns may suggest heteroscedasticity or model misspecification

**Fig. S8.**
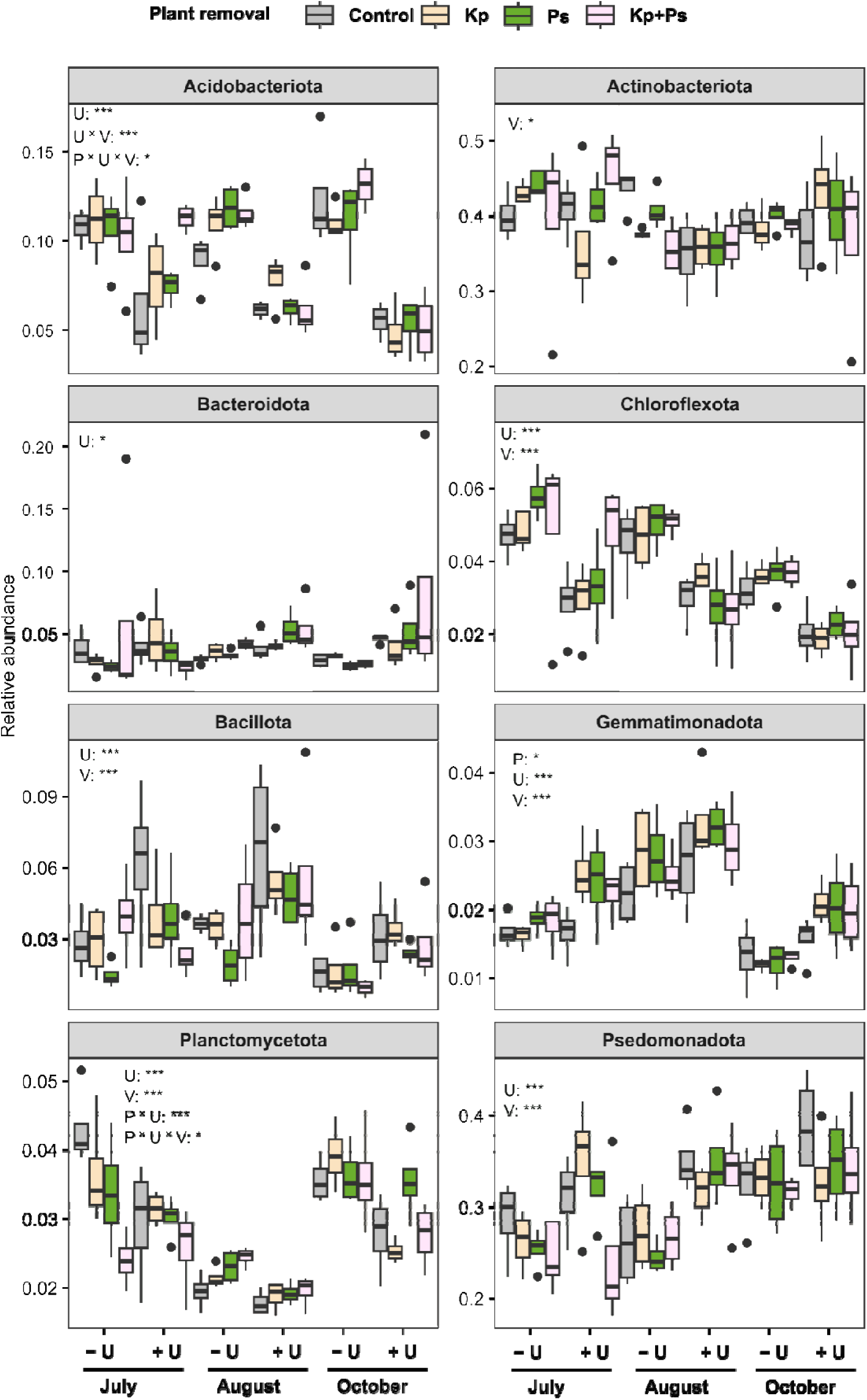
Effects of plant species removal and urea addition on the eight most abundant bacterial phyla across the growing season. Plant species removal treatments include no plant species removal (Control), removal of *Kobresia pygmaea* (Kp), removal of *Potentilla saundersiana* (Ps), and removal of both species (Kp+Ps). Urea treatments are represented as -U (without urea) and +U (with urea). Asterisks denote significant effects of dominant plant removal (P), urea addition (U), seasonal variation (V), and their interactions on the relative abundance of the respective phyla (\*\*\**p* < 0.001, \*\**p* < 0.01, \**p* < 0.05). Only significant results are displayed.

## Notes

### Competing Interest Statement

The authors have declared no competing interest.

